# PRDX6 contributes to selenocysteine metabolism and ferroptosis resistance

**DOI:** 10.1101/2024.06.04.597364

**Authors:** Zhiyi Chen, Alex Inague, Kamini Kaushal, Gholamreza Fazeli, Thamara N Xavier da Silva, Ancely Ferreira dos Santos, Tasneem Cheytan, Florencio Porto Freitas, Umut Yildiz, Lucas Gasparello Viviani, Rodrigo Santiago Lima, Mikaela Peglow Pinz, Isadora Medeiros, Thiago Geronimo Pires Alegria, Railmara Pereira da Silva, Larissa Regina Diniz, Simon Weinzweig, Judith Klein-Seetharaman, Andreas Trumpp, Adriana Mañas, Robert Hondal, Matthias Fischer, Christoph Bartenhagen, Briana K. Shimada, Lucia A. Seale, Marietta Fabiano, Ulrich Schweizer, Luis E. Netto, Flavia C. Meotti, Hamed Alborzinia, Sayuri Miyamoto, José Pedro Friedmann Angeli

## Abstract

Selenocysteine (Sec) metabolism is crucial for cellular function and ferroptosis prevention and has traditionally been thought to begin with the uptake of the Sec carrier selenoprotein P (SELENOP). Following uptake, Sec released from SELENOP undergoes metabolisation via selenocysteine lyase (SCLY), producing selenide, a substrate used by selenophosphate synthetase 2 (SEPHS2), which provides the essential selenium donor - selenophosphate - for the biosynthesis of the selenocysteine tRNA. Here, we report the discovery of an alternative pathway mediating Sec metabolisation that is independent of SCLY and mediated by peroxiredoxin 6 (PRDX6). Mechanistically, we demonstrate that PRDX6 can readily react with selenide and interact with SEPHS2, potentially acting as a selenium delivery system. Moreover, we demonstrate the presence and functional significance of this alternative route in cancer cells where we reveal a notable association between elevated expression of PRDX6 with a highly aggressive neuroblastoma subtype. Altogether, our study sheds light on a previously unrecognized aspect of Sec metabolism and its implications in ferroptosis, offering new avenues for therapeutic exploitation.

## Introduction

Ferroptosis is a non-appoptotic form of cell death characterized by an iron-dependent accumulation of phospholipid hydroperoxides, ultimately leading to membrane rupture and cell death (Dixon et al., 2012). Due to its association with various pathological conditions, including cancer, neurodegeneration, and tissue damage (Stockwell et al., 2017), the mechanisms regulating ferroptosis have garnered increasing interest. Moreover, ferroptosis has been put forward as a promissing strategy to eradicate therapeutically challenging entities such as neuroblastoma (Alborzinia et al., 2022), B-cell lymphoma (Freitas et al., 2024) and undifferentiated melanoma (Friedmann Angeli et al., 2019).

The selenoprotein glutathione peroxidase 4 (GPX4) is arguably the most important inhibitor of ferroptosis. GPX4 efficiently reduces phospholipid hydroperoxides to nontoxic lipid alcohols using glutathione (GSH) as a substrate (Friedmann Angeli et al., 2014; Ursini et al., 1982). Therefore, there is a growing interest to target this regulator as a means to elicit ferroptosis. Since GPX4 contains a selenocysteine (Sec) residue (U46) at its active site, responsible for its superior hydroperoxidase activity (Ingold et al., 2018), modulation of selenium/Sec metabolism is emerging as a prospective strategy to modulate ferroptosis (Alborzinia et al., 2023; Li et al., 2022). The synthesis of selenoproteins involves a well-understood multi-step process, including the biosynthesis of Sec-tRNA and the recoding of the UGA codon within the mRNA of selenoproteins (Labunskyy et al., 2014). Vital to fuel this process is the effective supply of selenium and its subsequent metabolization to the selenium donor selenophosphate (Burk and Hill, 2015). Thus, selenium supplementation is routinely employed as a strategy to elevate selenoprotein levels *in vitro* and *in vivo*. Conversely, restricting selenium uptake can broadly impair selenoprotein translation and increase ferroptosis sensitivity by of the loss of GPX4 (Alborzinia et al., 2023; Li et al., 2022). As such there is a growing interest in understanding and targeting Sec metabolism as a prospective strategy to induce ferroptosis(Dos Santos et al., 2023).

Cells in culture predominantly take up selenium through two pathways: one involving inorganic sources like selenite and the other supplying Sec primarily originating from selenoprotein P (SELENOP), a selenoprotein containing multiple Sec residues. Uptake of inorganic selenium in the form of selenite is promoted by system Xc- in a thiol dependent manner (Carlisle et al., 2020; Olm et al., 2009). Alternatively, the uptake of Sec-containing SELENOP requires its recognition primarily by the low-density lipoprotein receptor-related protein 8 (LRP8). Upon endocytosis, SELENOP is digested in the lysosome and free Sec in both reduced and oxidized form are made available for further metabolization. Notably, the low activity of system Xc^-^ in MYCN-amplified neuroblastoma disclosed an unforeseen dependency on LRP8, where the genetic deletion of this receptor was sufficient to induce ferroptosis (Alborzinia et al., 2023) indicating that the metabolism of Sec could be an actionable target in this entity. Of note, the lysosomal degradation of other selenoproteins can also provide cells with Sec accompanied by recycling of the Sec residue providing selenium to produce Sec-tRNA.

Direct recycling of Sec and incorporation into selenoproteins is not possible, because of the absence of a tRNA synthase charging tRNASec with Sec (Labunskyy et al., 2014). Specifically, serine, initially bound to tRNASec, is converted into Sec using selenophosphate. Hence, selenium must first be released from Sec and converted into selenophosphate. The essentail deselenation step is carried by selenocysteine β-Lyase (SCLY), which decomposes Sec to alanine and selenide (HSe^−^). SCLY is postulated to transfer HSe^−^ either directly to SEPHS2 in the form of a thioperselenide or to release it in the gaseous form (Tobe and Mihara, 2018). Once delivered, SEPHS2 phosphorylates HSe^−^ to generate H2SePO3^−^, the active form of Se required for the biosynthesis of the Sec-tRNA and the translation of new selenoproteins.

In the present study we demonstrate that, in proliferative cells, Sec-promoted selenoprotein biosynthesis can occur independently from SCLY, challenging the current paradigm. Using a loss of function CRISPR based screen, we uncovered that this parallel pathway utilizes peroxiredoxin 6 (PRDX6). Specifically, we show that PRDX6 can react with HSe^-^ and interact with SEPHS2, likely acting as an intracellular selenium carrier. Moreover, we demonstrate that high PRDX6 levels is associated with the pathology and relapse of neuroblastoma, suggesting that targeting this pathway could be exploited for therapeutic benefit in this entity.

## Results

### Identification of Sec-dependent regulators of ferroptosis

Previous works suggested that the utilization of Sec from SELENOP or the degradation of other selenoproteins entails the liberation of the selenium atom from Sec by SCLY (Seale et al., 2018) (Figure 1A). Given our recent identification of the essentiality of the SELENOP/LRP8 axis in *MYCN*-amplified neuroblastoma in preventing ferroptosis, we sought to investigate whether the absence of SCLY could mimic this observation and present an additional target to induce ferroptosis in this entity. To explore this, we employed the neuroblastoma cell line SK-N-DZ and engineered *SCLY*-deficient cells (*SCLY^KO^*) using three sgRNAs, with *LRP8* sgRNA serving as a control. *LRP8* deficient cells (*LRP8^KO^*) robustly reduced the expression of selenoproteins, demonstrated by the loss of selenoprotein N (SELENON) and GPX4, and induced ferroptosis as previously reported (Alborzinia et al., 2023). In contrast, *SCLY^KO^* did not affect the levels of these selenoproteins, while still triggering ferroptosis. However, cell death in *SCLY^KO^* could be prevented using lower concentrations of the ferroptosis inhibitor Liproxstatin-1 (Lip1) compared to *LRP8^KO^* cells, suggesting that the loss of SCLY does not fully phenocopy the loss of LRP8 (Figure 1B). Interestingly, supplementation of *SCLY^KO^* cells with L-selenocystine (the oxidized form of Sec) promoted a marked increase in the level of the selenoproteins GPX4 and SELENON (Figure 1C) and promoted their proliferation, suggesting the existence of a redundant pathway independent of SCLY in Sec metabolism. Therefore, we exploited this feature to identify factors that could be promoting the use of Sec in *SCLY^KO^*cell lines. In brief, we conducted a loss of function genome-wide CRISPR screen in a clonal *SCLY^KO^* SK-N-DZ cell line to identify alternative pathways that could contribute to the utilization of Sec in the absence of SCLY. The screen was designed with two conditions: L-selenocystine supplementation (50 nM) as the sole supplement preventing ferroptosis and fully-ferroptosis protective condition (500 nM Lip1 and 20 nM sodium selenite, Na2SeO3) (Figure 1D). The use of an alternative source of selenium in the Lip1 arm was deemed necessary to avoid differences in proliferation that could have arisen from the loss of TXNRD1, another selenoprotein shown to impact proliferation (Muri et al., 2018). Compared to the control condition, the screen identified a significant reduction in *PRDX6*-targeting sgRNAs under the L-selenocystine condition (Figure 1E), indicating that this protein could be involved in Sec utilization.

**Figure 1.**
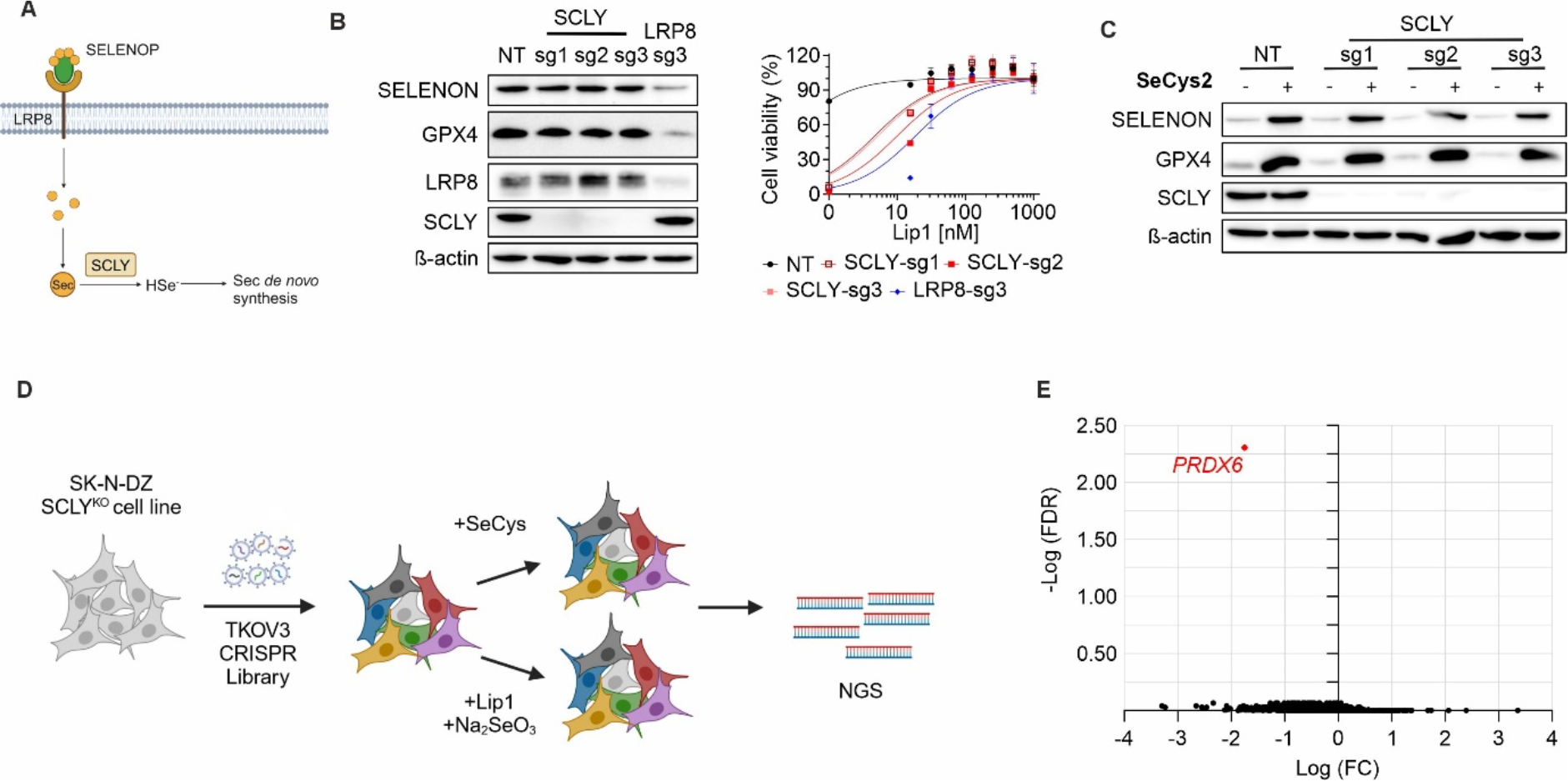
Identification of PRDX6 as a ferroptosis regulator A. Schematic representation of SCLY function promoting the release of selenium from Sec B. Generation and characterization of *SCLY* and *LRP8* deficient SK-N-DZ. Left panel, immunoblot analysis of SCLY, LRP8, GPX4 and SELENON expression in SK-N-DZ transduced with sgRNAs targeting *SCLY* or *LRP8*. Right panel, viability of cell lines, either wild-type or knock-out of *SCLY* or *LRP8* in the presence of increasing concentration of Lip1. Data are the mean ± SD of n=3 wells of 96-well plate. C. Immunoblot analysis of SCLY, SELENON and GPX4 expression in SK-N-DZ transduced with sgRNAs targeting *SCLY* with or without supplement of 50 nM L- selenocystine. D. Strategy of genome-wide CRISPR screen in SK-N-DZ *SCLY ^KO^* cell line. E. Result of CRISPR deletion screen displays -log FDR and log FC between 500 nM Lip1 + 10 nM Na2SeO3 and 50 nM Selenocystine.

### PRDX6 lipid hydroperoxidase activity does not contribute to ferroptosis resistance

Given the abundance of studies indicating that PRDX6, similar to GPX4, can function as a phospholipid hydroperoxidase (Chen et al., 2000), we initially reasoned that the screen might not have successfully identified Sec-specific effects. It seemed plausible that PRDX6 scored high due to the absence of this secondary phospholipid hydroperoxidase mechanism, a scenario unlikely to occur in the Lip1-treated arm of the screen (Figure 2A). To investigate whether the GPX4-like activity of PRDX6 contributes to the observed phenotype, we first revisited the reaction of rat PRDX6 (rPrdx6) against different hydroperoxides using stopped-flow spectroscopy, allowing us to determine rate constants of 3.0 ± 0.2 × 10^7^, 1.4 ± 0.04 × 10^5^ and 3.4 ± 0.1 × 10^4^ M^-1^s^-1^ for H2O2, LAOOH and PCOOH respectively (Figure S1A). Of note, rate constants with H202 are in agreement with previous reports (Toledo et al., 2011). Considering that peroxidized phospholipids are the primary substrates driving ferroptosis (Freitas et al., 2024), it is noteworthy that PRDX6 is approximately 3 to 4 orders of magnitude less efficient than GPX4 against these substrates, which has a reported rate constant close to 10^8^ M^-1^s^-1^ for PCOOH (Ursini et al., 2022). These results suggest that the phospholipid hydroperoxidase activity of PRDX6 is outcompeted by GPX4 *in vivo*. To demonstrate this, we initially overexpressed PRDX6, SCLY, LRP8, and FSP1 in Pfa1 cells (fibroblasts derived from the 4-hydroxytamoxifen (Tam)-inducible Gpx4^-/-^ model) (Seiler et al., 2008). This model allows for the genetic deletion of *Gpx4* upon treatment with Tam, leading to cell death. Of note, Pfa1 cells are characterized by low endogenous expression of *Lrp8* (Doll et al., 2017), and thus provide a controlled environment for studying the interplay between these factors (Figure 2B). Among the overexpressed cDNAs, only *FSP1* was able to complement *Gpx4* loss (Figure 2C- D). Importantly, PRDX6 was not able to complement genetic loss of *Gpx4*, arguing against a GPX4-like function. Moreover, only *LRP8* and *FSP1* overexpression could provide an increased resistance against ferroptosis inducers, rendering cells resistant to a panel of ferroptosis inducers (Figure 2E). To provide additional support for these observations, we employed an engineered cell line that lacks *GPX4* and relies exclusively on FSP1 for survival (HT1080 *GPX4*^KO^ *FSP1*-OE) (Doll et al., 2019)(Figure 2F and Figure S1B). Using these models, we can elicit ferroptosis by blocking FSP1 with the highly specific inhibitor iFSP1 (Nakamura et al., 2023; Xavier da Silva et al., 2023). Unlike the increased sensitivity of *PRDX6*, *SCLY* and *LRP8* deficient cells to canonical ferroptosis inducers, which act directly or indirectly on GPX4, genetic loss of any of those genes did not impact on ferroptosis sensitivity in a GPX4-independent context (Figure 2G and S1C), suggesting that PRDX6 is functionally upstream of GPX4.

**Figure 2.**
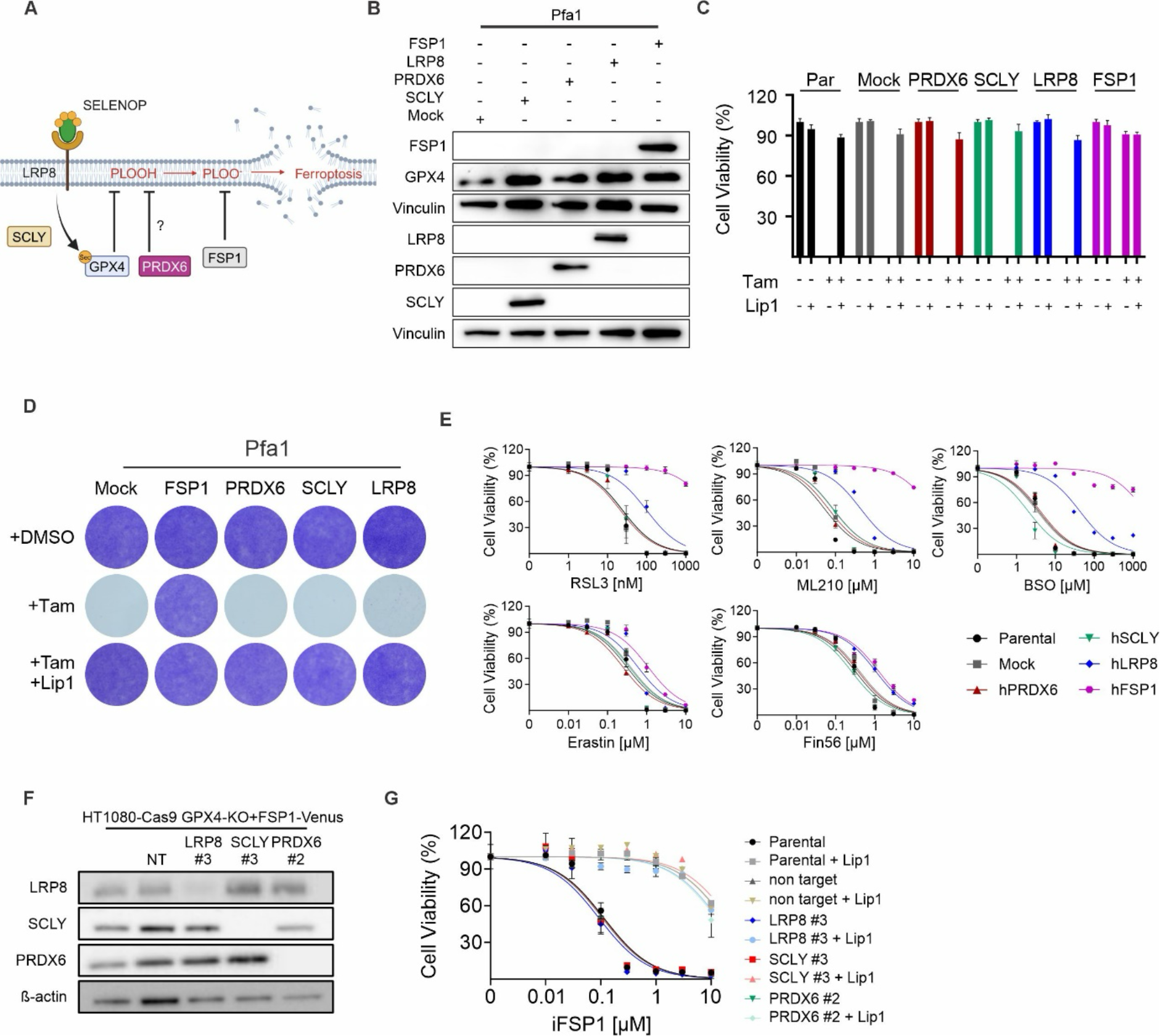
The lipid phospholipid hydroperoxidase activity of PRDX6 does not contribute to ferroptosis resistance A. Schematic representation of ferroptosis regulation positioning PRDX6 as a putative lipid hydroperoxidase. B. Immunoblot analysis of FSP1, SCLY, PRDX6, LRP8 and GPX4 expression in Pfa1 cell line over-expressing the indicated cDNAs. C. Viability of Pfa1 cell lines over-expressing the indicated cDNAs, in the presence of 500 nM tamoxifen or 500 nM Lip1. Par=parental cell line. Data are the mean ± SD of n=3 wells of 96-well plate. D. Clonogenic capacity of Pfa1 cell lines over-expressing the indicated cDNAs in the presence of DMSO, tamoxifen (500nM) or tamoxifen and Lip1 (500 nM). E. Viability of Pfa1 cell lines over-expressing the indicated cDNAs in the presence of increasing concentrations of RSL3, Fin56, Erastin and BSO. Data are the mean ± SD of n=3 wells of 96-well plate. F. Immunoblot analysis of SCLY, PRDX6 and LRP8 expression in HT1080-Cas9 with GPX4^KO^ + FSP1^oE^ cells transduced with sgRNAs targeting LRP8, PRDX6 and SCLY. G. Viability of cell lines depicted in (F), challenged with increasing concentration of iFSP1 in the presence or absence of Lip1 (500 nM). Data are the mean ± SD of n=3 wells of 96-well plate.

### PRDX6 modulates Sec metabolism and ferroptosis sensitivity

Having formally excluded a potential GPX4-like function of PRDX6 in ferroptosis , we next set to functionally explore PRDX6 potential role in Sec metabolism. For this, we deleted *PRDX6* using three sgRNAs in the SK-N-DZ cell line (*PRDX6^KO^*). *PRDX6* loss resulted in ferroptosis, albeit *PRDX6^KO^* cells could be rescued with lower concentrations of Lip1 compared to *LRP8^KO^* cells (Figure 3A, right). Additionally, *PRDX6^KO^* cells were characterised by decreased expression levels of SELENON and GPX4, resembling LRP8 deficiency (Figure 3A, left). Treatment with L-selenocystine (50nM) induced an increase in GPX4 and SELENON levels in the *PRDX6^KO^* cell lines, albeit to a lesser extent than in the wild-type cell line (Figure 3B), indicating a role for PRDX6 in the metabolism of Sec. This assumption was further reinforced by analysing LRP8 co-dependencies using the DepMap (www.depmap.org) (Figure 3C). This computational approach builds on the observation that genetic perturbations affecting components within the same or related pathways often exhibit similar dependencies. As expected, LRP8 co-dependency analysis revealed several enzymes implicated in selenoprotein biosynthesis (EEFSEC, SCLY, SEPHS2, and SEPSECS). Notably, PRDX6 scored as the highest co-dependency with LRP8, thus strengthening a functional relationship between these two proteins and the metabolism of Sec (Fig. 3C) in line with previous computational work (Santesmasses and Gladyshev, 2022).

**Figure 3.**
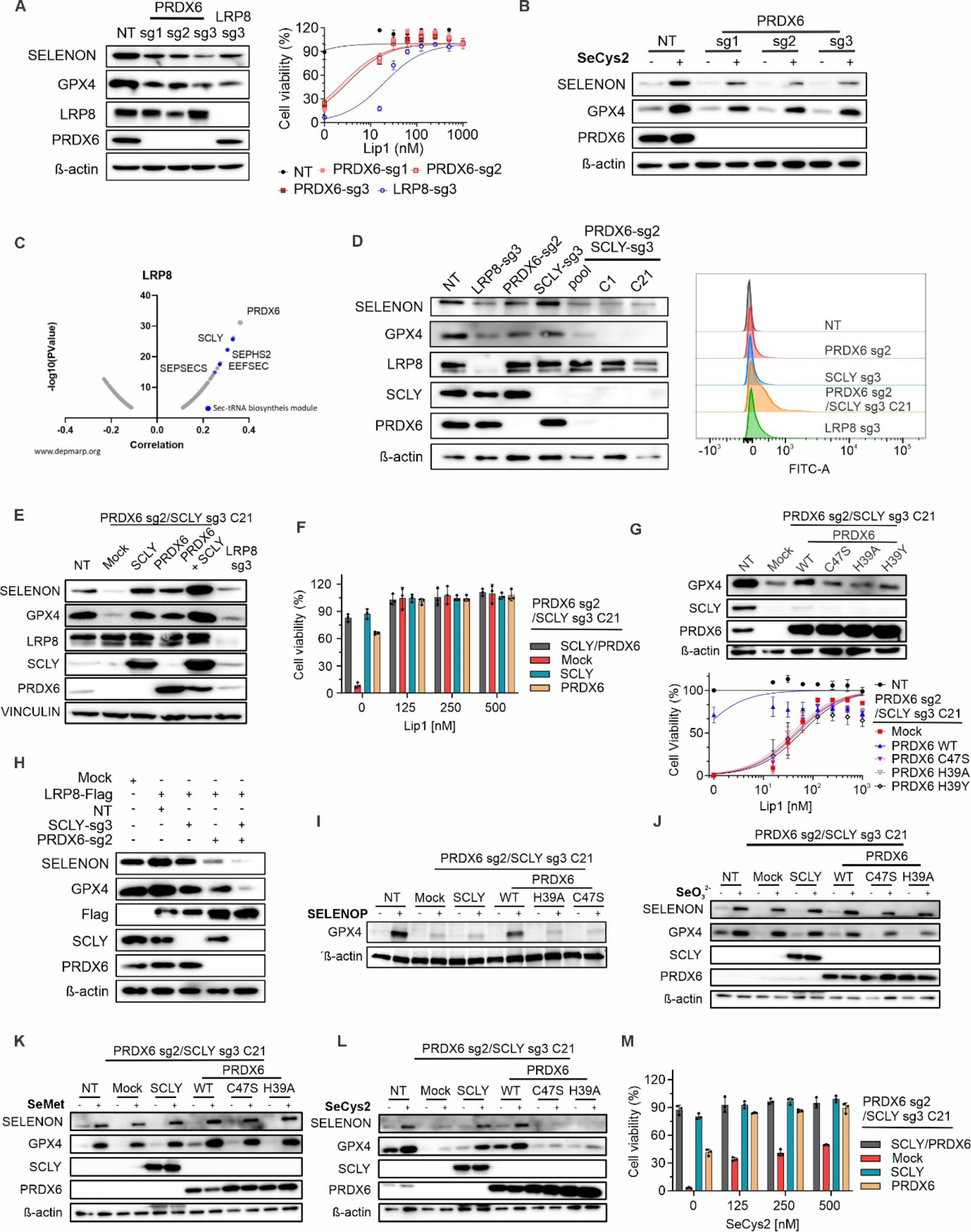
PRDX6 modulates Sec metabolism and ferroptosis sensitivity A. Generation and characterization of *PRDX6* and *LRP8* deficient SK-N-DZ. Left panel, immunoblot analysis of PRDX6, LRP8, GPX4 and SELENON expression in SK-N-DZ transduced with sgRNAs targeting *PRDX6* or *LRP8*. Right panel, viability of cell lines, either wild-type or knock-out of *PRDX6* and *LRP8* in the present of increasing concentration of Lip1. Data are the mean ± SD of n=3 wells of 96-well plate. B. Immunoblot analysis of PRDX6, SELENON and GPX4 expression in SK-N-DZ transduced with sgRNAs targeting *PRDX6* with or without supplement of 50 nM L- selenocystine. C. Co-dependency analysis demonstrating the high degree of co-essentiality between LRP8 and PRDX6 (www.depmap.org). D. Left panel, immunoblot analysis of LRP8, SELENON, SCLY, PRDX6 and GPX4 expression in SK-N-DZ transduced with sgRNAs targeting *LRP8*, *PRDX6* and *SCLY.* C1: monoclonal 1; C21: monoclonal 21. Right panel, flow cytometry analysis of BODIPY 581/591 C11 oxidation in SK-N-DZ transduced with sgRNA against *LRP8, PRDX6* and *SCLY*, after Lip1 removal for 24 hours. E. Immunoblot analysis of LRP8, SELENON, SCLY, PRDX6 and GPX4 in the wild-type SK-N-DZ, *LRP8* deficient cells and a monoclonal *PRDX6* and *SCLY* deficient SK-N- DZ with SCLY and PRDX6 restoration. F. Viability of cell lines, either wild-type or knock-out of *PRDX6*, knock-out of *SCLY* and knock-out of *PRDX6* and *SCLY* in the present of increasing concentration of Lip1. Data are the mean ± SD of n=3 wells of 96-well plate. G. Immunoblot analysis of LRP8, SELENON, SCLY, PRDX6 and GPX4 in the wild-type SK-N-DZ and monoclonal PRDX6 and SCLY deficient SK-N-DZ with PRDX6 wild-type (WT), C47S, H39A and H39Y restoration. Lower panel shows the viability of the cell lines in the present of increasing concentration of Lip1. Data are the mean ± SEM of n=3 wells of 96-well plate from three independent experiments. H. Immunoblot analysis of Flag, SELENON, SCLY, PRDX6 and GPX4 expression in wild- type SK-N-DZ and SK-N-DZ overexpressed LRP8-Flag transduced with sgRNAs targeting *PRDX6* and *SCLY*. I. Immunoblot analysis of GPX4 in the wild-type and monoclonal *PRDX6/SCLY*-deficient SK-N-DZ with PRDX6 WT, C47S, H39A and SCLY restoration with supplement of SELENOP. J. Immunoblot analysis of SELENON, SCLY, PRDX6 and GPX4 in the wild-type and monoclonal PRDX6/SCLY deficient SK-N-DZ with PRDX6 wild-type, C47S, H39A and SCLY restoration with supplement of 50 nM Na2SeO3. K. Left panel, immunoblot analysis of SELENON, SCLY, PRDX6 and GPX4 in the wild- type and monoclonal *PRDX6/SCLY* deficient SK-N-DZ with PRDX6 wild-type, C47S, H39A and SCLY restoration with supplement of 5 µM L-Selenomethione. L. Immunoblot analysis of SELENON, SCLY, PRDX6 and GPX4 in the wild-type and monoclonal PRDX6/SCLY deficient SK-N-DZ with PRDX6 wild-type, C47S, H39A and SCLY restoration with supplement of 50 nM L-selenocystine. M. Viability of cell lines, either wild-type or knock-out of *PRDX6*, knock-out of *SCLY* and knock-out of *PRDX6* and *SCLY* in the present of increasing concentration of L- selenocystine. Data are the mean ± SD of n=3 wells of 96-well plate.

Given that neither PRDX6 or SCLY loss fully recapitulated the loss of LRP8, and both could still utilize Sec for the biosynthesis of new selenoproteins, we explored a potential redundancy between SCLY and PRDX6. Therefore, we generated *SCLY* and *PRDX6* double deficient cells (*SCLY^KO^/PRDX6^KO^*). Similar to *LRP8^KO^* and unlike the single deletion of SCLY or PRDX6, knockout of both genes strongly decreased GPX4 and SELENON expression (Figure 3D, left). We next assessed the propensity to spontaneously undergo lipid peroxidation upon Lip1 withdrawal in these cell lines. Following Lip1 removal for 24 hours, lipid peroxidation was assessed using BODIPY-C11, revealing that *SCLY^KO^/PRDX6^KO^* resulted in a stronger increase in lipid peroxidation compared to *LRP8^KO^*and the single deficiencies of either *SCLY* or *PRDX6* (Figure 3D, right). To expand on this observation, we deleted SCLY, and PRDX6 alone and in combination in a panel of cell lines, reaffirming that *SCLY^KO^* and *PRDX6^KO^* disrupted GPX4 expression, with the *SCLY^KO^/PRDX6^KO^*showing a more pronounced loss of selenoproteins (Figure S2A). Notably, in non NB cell lines neither single deficiency (*SCLY^KO^* or *PRDX6^KO^*) nor double deficiency (*SCLY^KO^/PRDX6^KO^*) resulted in ferroptosis, a phenotype similar to what we recently reported for the loss of LRP8 (Alborzinia et al., 2023). Nonetheless, they showed increased sensitivity to a set of ferroptosis-inducing compounds (Figure S2B) and were also associated to an increased basal level of BODIPY-C11 oxidation, a surrogate of increased lipid-peroxidation (Figure S2C).

To provide additional validation into the function of PRDX6 in modulating selenoprotein expression, we reintroduced *PRDX6* and *SCLY* in the *SCLY^KO^/PRDX6^KO^* SK-N-DZ (C21) cells where we could observe a rescue of GPX4 and SELENON expression (Figure 3E) upon the reconstitution of *PRDX6* or *SCLY*. Moreover, the forced expression of either *PRDX6* and *SCLY* partially rescued the double knock-out cells from ferroptosis, suggesting that the proteins act in parallel through non-overlapping mechanisms (Figure 3F). We next focused on identifying features that promote Sec metabolization by PRDX6. A set of PRDX6 mutants lacking peroxidase activity (C47S) and mutants where the sulfenic acid is destabilized (H39A or H39Y)(Lang et al., 2023) was generated and studied for their impact in modulating selenoprotein levels. Our results showed that in the *SCLY^KO^/PRDX6^KO^*, only restoration of wild-type PRDX6 (WT), but not C47S, H39A or H39Y variants rescued GPX4 expression and cell viability (Figure 3G). These data highlight the critical importance of the catalytic cysteine residue and the proper configuration of the PRDX6 active site in promoting selenoprotein expression and in protection against ferroptosis.

Identified the high degree of co-dependency between PRDX6 and LRP8 (Figure 3C), we reasoned that PRDX6 may operate downstream of LRP8. To investigate this, we generated *PRDX6^KO^*, *SCLY^KO^*and the *SCLY^KO^/PRDX6^KO^* in SK-N-DZ cells overexpressing LRP8. Using these cell lines, we revealed that the increase in selenoprotein production stimulated by the increased uptake of SELENOP in *LRP8*-overexpressing cells is disrupted in *PRDX6^KO^* and completely abrogated in the *SCLY^KO^/PRDX6^KO^* (Figure 3H). The supplementation with radioactively labelled SELENOP (^75^Se-SELENOP) confirmed the impaired incorporation of ^75^Se in (Figure S3A). Recapitulating the previous observations, only the reconstitution of PRDX6 WT, but not C47S or H39A mutants, was capable of rescuing GPX4 expression in *SCLY^KO^/PRDX6^KO^* cells upon SELENOP supplementation (Figure 3I). To test whether PRDX6 is selective towards the type of selenium source it metabolises, we treated the add-back model with L-selenocystine, L-selenomethionine and sodium selenite (Na2SeO3). Interestingly, Na2SeO3 (Figure 3J) and L-selenomethionine (Figure 3K) could equally promote the expression of GPX4 and SELENON. However, Sec-induced translation of selenoproteins in the *SCLY^KO^/PRDX6^KO^* cells was strongly impaired (Figure 3L), suggesting that the effect of PRDX6 is associated with the metabolism of Sec and it’s not limited to SELENOP. In the add-back model, administrating increasing concentration of L-selenocystine efficiently rescued the cells with reintroduced PRDX6 and SCLY (Figure 3M). To evade the artifact of the forced expression, L-selenocystine and Na2SeO3 were also applied in the *PRDX6^KO^*, *SCLY^KO^*, with endogenous level of PRDX6 and SCLY, and the *SCLY^KO^/PRDX6^KO^* cells. L-selenocystine and Na2SeO3 could both partially rescue the viability of *SCLY^KO^*and *PRDX6^KO^* to a similar extent, while, in *SCLY^KO^/PRDX6^KO^*, L-selenocystine was less efficient (Figure S3B).

Furthermore, we confirmed that PRDX6 regulates selenoprotein expression via selenocysteine metabolism in another neuroblastoma cell line, NMB, supplemented with SELENOP (Figure S3C), L-selenocystine (Figure S3D), L-selenomethionine (Figure S3E) and Na2SeO3 (Figure S3F). Altogether, these data demonstrate that PRDX6 is preferentially required for the metabolization of organic selenium in the form of Sec and that this is not limited to the Sec arising from exogenous SELENOP.

### PRDX6 reacts with hydrogen selenide

After demonstrating that PRDX6 promotes selenoprotein biosynthesis, a mechanistic link was still missing. Initial attempts did not show that PRDX6 has a deselenase activity, making it unlikely that it acts similarly to SCLY. PRDX6 is notable for existing as a stable sulfenic acid (Akter et al., 2018; Choi et al., 1998; Kim et al., 2016). This feature could promote its interaction with HSe^-^, similar to reactions between sulfenic acids found in related peroxiredoxins and H2S (Cuevasanta et al., 2019). This reaction could lead to the formation of a thioperselenide intermediate that mediates the transfer of selenium to SEPHS2.

To provide evidence for this mechanism, we investigated whether PRDX6 can react directly with different selenium sources. We studied the reaction products of rPrdx6 with L-selenocystine, Na2SeO3, and Na2Se using non-digested protein analysis by mass spectrometry (Figure 4A-C and S4A). Remarkably, we only detected the formation of a new product at pH 7.4 when Na2Se was used as a source of selenium. In aqueous solution, Na2Se readily produces the gas H2Se. The experiment detected a predominant mass shift indicating the formation of a rPrdx6-S-SeO2H product (Figure 4A). The identity of the product was confirmed by incubating the PRDX6 (C47S) mutant with Na2Se, where the 112-mass shift corresponding to SeO2H was completely eliminated. The results are in line with the formation of a thioperselenide, which is produced by the reaction of HSe^-^ and the sulfenic acid of PRDX6. This thioperselenide then undergoes further oxidation during MS analysis (see Figure 4C) producing the characteristic thioselenic acid (S-SeO2H).

**Figure 4.**
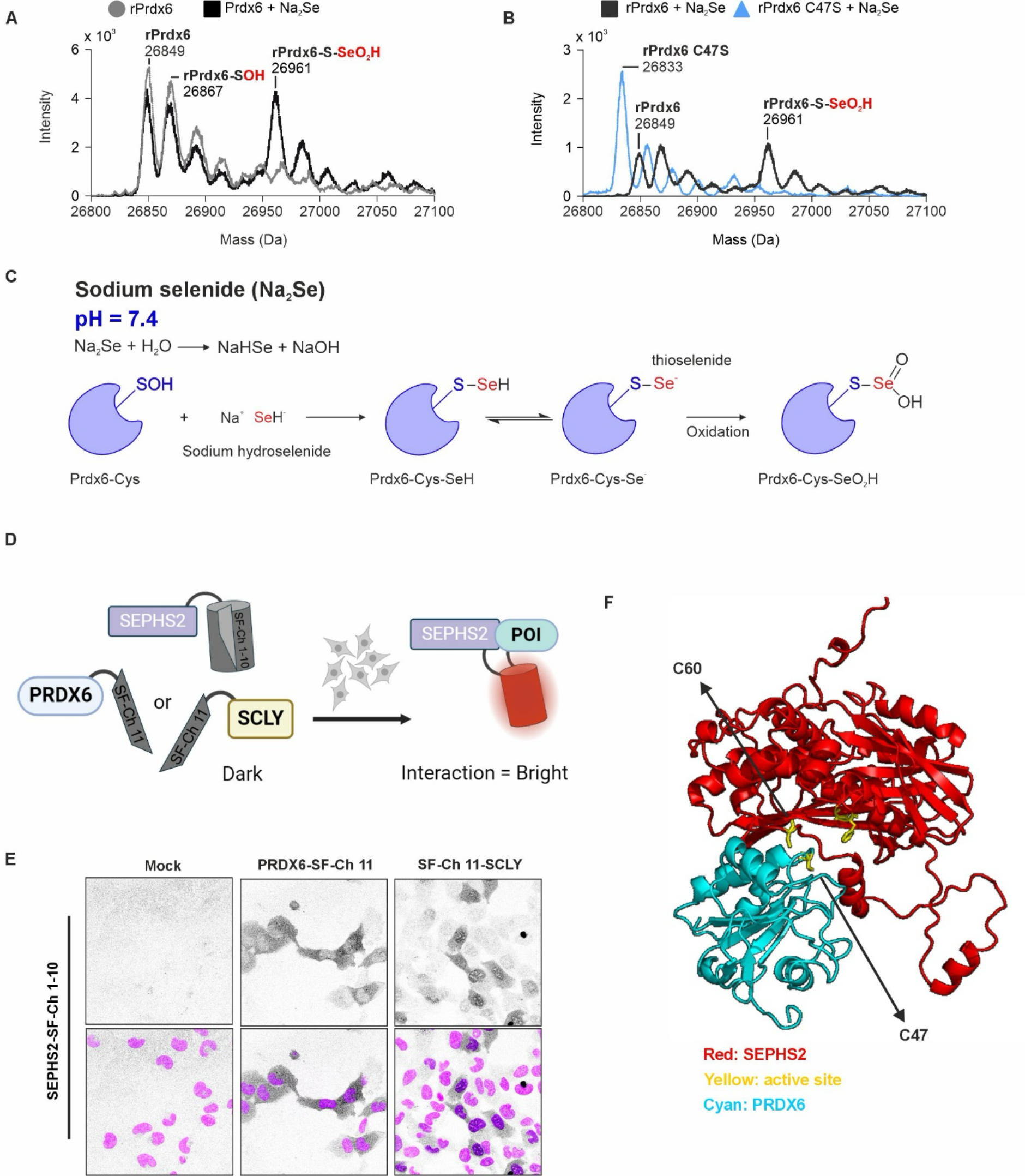
PRDX6 links HSe^-^ to SEPHS2 A. Mass spectra obtained from the incubation of 35 µM rPrdx6 with 1 mM Na2Se. (pH 7.4, 500 rpm, 37 °C). B. Mass spectra obtained from incubation of 35 µM rPrdx6 or rPrdx6 C47S with 1 mM Na2Se. (pH 7.4, 500 rpm, 37 °C). C. Schematic representation of the putative reaction between rPrdx6 and HSe produced from Na2Se decomposition. D. Schematic representation of the split fluorescent protein strategy to assess the interaction of SEPHS2 with PRDX6 or SCLY (see also Fig. S6A-D). E. Representative images of cells transfected with SEPHS2-SF-Ch 1-10 controls (left), together with PRDX6-SF-Ch 11 (middle), or SF-Ch-11-SCLY (right). Dark tones show the formation of fluorescent SF-Ch. Hoechst staining (magenta) is added in the lower panels to mark the nuclei. F. Docking analysis of the potential interaction between SEPHS2 (red) with PRDX6 (cyan). Active sites of proteins are indicated in yellow. Potential critical amino acids are marked.

Upon incubating rPrdx6 with L-selenocystine, a mass increase of approximately 167 Da was detected in the protein at pH 5.0, while no such increase was observed at pH 7.4 (Figure S4B). This addition aligns with the incorporation of a L-selenocysteine (Figure S4A). Similarly, an increase of 112 Da was exclusively observed when incubating rPRDX6 with Na2SeO3 at pH 5.0, with no corresponding change at pH 7.4 (Figure S4C). This indicates the formation of a SeO2H adduct. As anticipated, no reaction was observed with L-selenomethionine (Figure S4D), wherein the selenium atom is shielded by a methyl group. In all cases, none of the products were detected with the rPrdx6 C47S mutant (Figure S4E), further indicating that the peroxidatic cysteine of PRDX6 reacts with selenium compounds. Taken together, these findings suggest that PRDX6 is capable of engaging various selenium compounds under acidic pH conditions, while exhibiting selectivity towards small molecules such as HSe^-^ under neutral (cytosolic) conditions. To better understand the observed selectivity at pH 7.4 (Figure 4C and S4F) for HSe^-^ we performed molecular dynamic (MD) simulation of rPrdx6. We first analyzed the changes in the solvent accessible surface area (SASA) of rPrdx6 active site along the simulations. Remarkably, the SASA values of the active site are overall higher among the simulations performed at pH 5 (mean = 521.5 ± 52.2 Å²; median = 520.2 Å²) than at pH 7 (mean = 407.3 ± 54.5 Å²; median = 407.7 Å²) (Figure S4G). Additionally, the average SASA values of the active site at pH 5 remained higher over time, compared to pH 7 (Figure S4H). We then analysed the changes in the SASA value of the catalytic residue C47 individually. Similar to what is observed for the active site as a whole, the SASA values of C47 were higher among the simulations at pH 5 (mean = 24.6 ± 8.8 Å²; median = 25.2 Å²) than at pH 7 (mean = 7.4 ± 6.3 Å²; median = 7.4 Å²) (Figure S4I), and the average SASA values of C47 at pH = 5 remained higher over time compared to pH 7 (Figure S4J). Together, these data indicate that the rPrdx6 active site, including the catalytic residue C47 and the corresponding sulfenic acid, is less solvent-exposed at pH 7. This suggests that only small and diffusible selenium compounds, such as HSe^-^, would have access to the active site potentially also preserving the stability of the thioperselenide intermediate.

The identification of a thioperselenide intermediate is important as it has been proposed that this could be the intermediate donating selenium to SEPHS2 (Lacourciere et al., 2002). Thus, we investigated a potential interaction between PRDX6 and SEPHS2 that could be critical to facilitate the transfer of the selenium atom between the two enzymes. For this purpose, we fused split parts of super-folder Cherry to these proteins (and SCLY). We tagged N- or C-terminus of SEPHS2U60C with SF-Ch-1-10, while SF-Ch11 labelled either terminus of PRDX6 or SCLY (Figure S5A). The constructs were stably transduced into HT1080 cells in various combinations (Figure S5B). Fluorescent signal was detected in the combination of C-terminal tagged PRDX6 (PRDX6-SF-Ch 11) and C-terminal tagged SEPHS2 U60C (SEPHS2-SF-Ch 1-10), and the combination of N-terminal tagged SCLY (SF-Ch 11-SCLY) and SEPHS2-SF- Ch 1-10, indicating the interaction of SEPHS2 with PRDX6 and with SCLY respectively (Figure S5C and 4D-E). We validated the functionality of tagged PRDX6 by demonstrating that it could restore GPX4 and SELENON expression as well as cell viability, by expressing it in *PRDX6^KO^* SK-N-DZ (Figure S5D). Duolink-PLA in *PRDX6* wild-type (*PRDX6^WT^*) and *PRDX6^KO^* SK-N-DZ were used as an orthogonal approach to demonstrate the close interaction of PRDX6 and SEPHS2 in cells, (Figure S5E). Given difficulties to establish a robust SEPHS2 enzymatic assay due to the fact that the protein itself is a selenoprotein, we explored further the potential interaction between PRDX6 and SEPHS2. The pairing of PRDX6 and SEPHS2 was used as input into ClusPro with each resulting in up to 30 different poses. The distance between active sites for every pose from PRDX6 and SEPHS2 was measured. The shortest distances between active sites were then visualized through PyMOL and the different poses were then compared. Across all PRDX6 and SEPHS2 combinations, the shortest distances for each combination were all below 15 angstroms with the shortest distance being 9.8 angstroms (Figure 4F), providing a model for the close interaction between their active sites.

One aspect that remains unresolved is how Sec is deselenated to produce HSe^-^. Given the chemical similarity between Sec and cysteine (Figure S4A), we conducted experiments to explore whether H2S biosynthesis pathway (Figure S5F) could promote HSe- production from Sec and act upstream of PRDX6. Using an SCLY^KO^ deficient cell line (Figure S5G) we generated cell lines deficient for CTH, CBS and MPST and posited that if they would produce HSe^-^ they should phenocopy the deficiency in selenoprotein production stimulated by selenocystine as observed for the dual loss of SCLY/PRDX6. Nevertheless, the single loss of any of the H2S generating enzyme did not influence the metabolization of L-selenocystine (Figure S5H-J). These results suggest that there are redundancies between the different H2S producing pathways or non-enzymatic mechanisms might be contributing to Sec deselenation.

### PRDX6 contribution to neuroblastoma growth

Our data show that PRDX6 and SCLY regulate selenoprotein production and ferroptosis by modulating Sec metabolism. Given the reported importance of this pathway for neuroblastoma growth we next evaluate its contribution in neuroblastoma models. Subsequently, we orthotopically implanted the SK-N-DZ LRP8^OE^ cell line with PRDX6/SCLY-proficient (*PRDX6^WT^/SCLY^WT^*), *SCLY^KO^*, *PRDX6^KO^* or *PRDX6^KO^/ SCLY^KO^* (Figure 3H) into the adrenal gland of NOD.Cg - Prkdc^scid^Il2rgtm1^Wjl^/SzJ (NSG) mice (Figure 5A). Compared with *PRDX6^WT^/SCLY^WT^*, *SCLY^KO^*and *PRDX6^KO^* showed a reduction in tumor growth, and the *PRDX6^KO^/SCLY^KO^*had a stronger effect on the LRP8 promoted growth in these xenografts (Figure 5B). Consistent with the *in vitro* findings, GPX4 levels were decreased in *PRDX6^KO^*tumors and the *SCLY^KO^* had only limited effect on GPX4, while a robust loss of GPX4 protein occurred in *PRDX6^KO^/SCLY^KO^* tumors (Figure 5C). Considering the strong impact of *PRDX6^KO^/SCLY^KO^* on tumor growth and GPX4 expression, we further explored the potential role of PRDX6 and SCLY in the establishment and growth of the tumor. Hence, we performed a follow-up experiment by implanting the SK-N-DZ with *PRDX6^WT^/SCLY^WT^*and *PRDX6^KO^/SCLY^KO^* in the adrenal gland of NSG mice (Figure S6A). To prevent ferroptosis, both cell lines were grown in the presence of Lip1 before implantation. Subsequently, in contrast to *PRDX6^WT^/SCLY^WT^*cells, *PRDX6^KO^/SCLY^KO^* cells exhibited a remarkable effect on tumor growth, directly impacting the overall survival of mice, with a median survival of 36 days in the KO group compared to 26 days in the WT group (Figure S6B).

**Figure 5.**
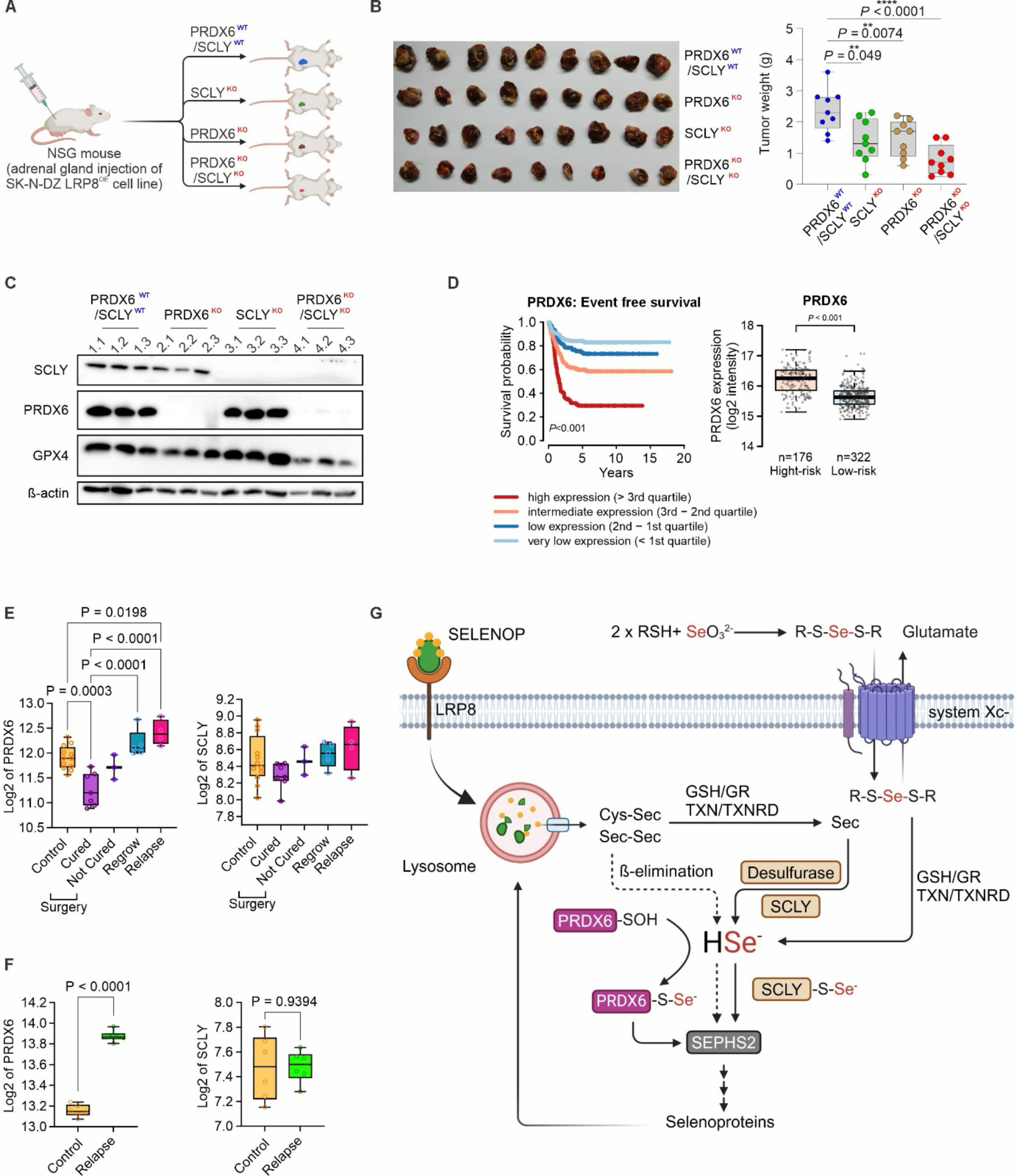
Sec metabolism contribute to neuroblastoma growth A. Schematic representation of the orthotopic implantation of control (*PRDX6^WT^/SCLY^WT^*) or *PRDX6*-deficient (*PRDX6^KO^/SCLY^WT^*) or *SCLY*-deficient (*PRDX6^WT^/SCLY^KO^*). B. Left panel, representative images of implanted orthotopic xenografts. Right panel, Tumor weight of orthotopically implanted control (*PRDX6^WT^/SCLY^WT^*; blue, n=9), *SCLY*-deficient (*PRDX6^WT^/SCLY^KO^*; green, n=9), *PRDX6*-deficient (*PRDX6^KO^/SCLY^WT^*; red, n=9) and *PRDX6/SCLY*-deficient (*PRDX6^KO^/SCLY^KO^*; purple, n=9). C. Immunoblot analysis of SCLY, PRDX6 and GPX4 levels from orthotopic tumors of *PRDX6^WT^/SCLY^WT^*, *PRDX6^WT^/SCLY^KO^*, *PRDX6^WT^/SCLY^WT^* and *PRDX6^KO^/SCLY^KO^* SK-N-DZ LRP8^OE^ cell lines. D. Association between PRDX6 on KaplanMeier survival analysis and expression levels (log2 intensity) in low-risk (LR) and high-risk (HR) neuroblastoma patients (n=459). The p-values were calculated using a two-sided Wilcoxon rank-sum test (boxplots, comparison of gene expression between high and low-risk patients) and log-rank test (Kaplan-Meier curves, pairwise comparisons) and Benjamini-Hochberg corrected. All p-values were adjusted for multiple testing (Benjamini- Hochberg). The centreal band indicates the median. The boxes cover the first and third quartile, with whiskers indicating the minimum and maximum of the data within 1.5× the interquartile range. E. Expression of *PRDX6* and *SCLY* across the COJEC treatment group in responsive PDX (PDX3). F. Expression of *PRDX6* and *SCLY* in organoids derived from responsive and relapsed PDX3. G. Schematic representation of the mechanism describing PRDX6 role in Sec metabolism

We next explored whether the PRDX6/SEPHS2 axis could be a predictor of clinical outcome in cohorts of pediatric neuroblastoma patients and in preclinical models. The high PRDX6 expression presented a strong correlation with poor survival outcomes (Figure 5D). Interestingly, we could extrapolate these observations in the clinically relevant patient-derived xenograft (PDX) models described in Mañas et al.,(Manas et al., 2022) where a chemotherapy regimen resembling COJEC, incorporating all five COJEC drugs, was used to treat neuroblastoma PDX models mirroring the characteristics of neuroblastoma patient tumors (Figure S6C). Re-analysis of this dataset using data from two PDX models allowed us to identify that PRDX6 expression is increased in COJEC refractory tumors. Moreover, clinical outcome in the PDX that was responsive to COJEC followed by surgical intervention demonstrated that high PRDX6 expression was associated with relapsing tumors, while lower expression was associated with curative response (Figure 5E-F and S6D-E). Taken together, these findings suggest that PRDX6 can impact selenium usage in patient-derived models and that the expression and likely activity of this module appear to link to disease progression and resistance to clinically relevant regiments.

## Discussion

It has long been suggested that the utilization of Sec, arising from the breakdown of SELENOP and other selenoproteins would depend exclusively on the activity of SCLY to release the selenium atom that would be subsequently channeled to produce selenophosphate. However, the relatively mild phenotype of mice deficient in *Scly* remained an unexplained puzzle (Seale et al., 2012). Previous works from Thressa Stadtmanńs, the discoverer of Sec, group in the early 2000s have argued that cells would require HSe- shuttling system given that the direct use of HSe- as a substrate for the reaction carried out by SEPHS2 is within a concentration that is toxic to cells (Kim et al., 1997; Lacourciere et al., 2002; Lacourciere et al., 2000). One of such delivery systems has been proposed to depend on SCLY, which promotes the deselenation step. Nevertheless, mouse studies have demonstrated that lack of SELENOP is not phenocopied by the loss of SCLY (Mizuno et al., 2023), indicating that the Sec derived from SELENOP can still be used for the production of new selenoproteins in the absence of SCLY. Moreover, the transport of SELENOP can be independent of SCLY according to the proliferative cell type, but the mechanism or the identity of this delivery system after SELENOP breakdown has remained largely uncharacterized. Here we report on an important step into the characterization of these SCLY-independent pathways, where an unforeseen contribution of PRDX6 in Sec metabolism and ferroptosis resistance is disclosed. Importantly, our work shows that loss of PRDX6 sensitizes cells and promotes ferroptosis via a non-canonical mechanism independent of its more well-established lipid hydroperoxidase activity.

Our current understanding based on the chemical reaction of PRDX6 with HSe^-^ and its close interaction with SEPHS2 is that PRDX6 could be acting to promote the productive use of HSe^-^. While we explored the kinetics of the reaction of PRDX6 with HSe^-^, we were unable to precise the products of the reaction based on the intrinsic protein fluorescence changes and hence, we could not specify the rate constants. Importantly, the use of Sec for selenoprotein production indispensably requires a desselenation step, and while this activity is well described for SCLY, it remains to be identified which enzymes are involved in the production of HSe- in its absence. Our initial attempts to identify the enzyme responsible for this focused on the known proteins able to produce the closely related gas H2S, namely CBS, CTH and MPST (Barayeu et al., 2023). We show that single deficiency of any of these three H2S-producing proteins did not impair Sec mobilization when SCLY was absent. We therefore hypothesize that HSe^-^ production from Sec is entailed by redundant activities between CBS, CTH and MPST or catalysed by promiscuous desulfurases, for example NFS1 which has been associated with ferroptosis resistance(Alvarez et al., 2017). Nonetheless, the enzymatic deselenation by desulfurases would be in direct competition with the metabolization of cysteine, rendering it greatly ineffective as cysteine concentrations within mammalian cells is believed to be three to four orders of magnitude higher than Sec. However, ineffective bacterial and archaeal desulfurases have been shown to carry SCLY-like reaction in the presence of cysteine (Mihara et al., 1997), therefore this possibility should not be formally excluded. Another possibility is that selenocystine could undergo spontaneous ß-elimination in cells to generate Sec, alanine and HSe-. The desselenation of selenoproteins is a well-studied phenomenon and entails the oxidation of the diselenide leading to deselenation of the Sec residue. The diselenide can be oxidized to R-Se-Se(O2)-R, followed by hydrolysis to Sec and Sec-seleninic acid which can rapidly undergo ß-elimination (Mousa et al., 2017). Further studies will be required to fill this knowledge gap (Figure 5G).

Moreover, motivated by the described dependency of neuroblastoma on selenocysteine uptake we investigated whether loss of SCLY and PRDX6 would impair growth under conditions of increased SELENOP uptake (achieved here by overexpressing LRP8) in an *in vivo* orthotopic model. Using this model, we demonstrate that deletion of SCLY or PRDX6 impair the growth of this xenograft and that a more pronounced growth inhibition was achieved by the dual loss of SCLY and PRDX6. Lastly, analyzing a large cohort of human patients and a PDX-relevant therapy model we could demonstrate that increased expression of PRDX6/SEPHS2 are associated with poor prognosis and also relapse.

Taken together Our study provides a fundamental contribution into the understanding of Sec metabolism and selenoprotein biosynthesis by establishes a new function for PRDX6 in promoting the productive use of HSe- in an SCLY-independent manner. Moreover, our study strongly suggests that the PRDX6/SEPHS2 is an actionable target that could be used to sensitize neuroblastoma to ferroptosis.

## Acknowledgments

J.P.F.A. acknowledges the support of the Junior Group Leader program of the Rudolf Virchow Center, University of Würzburg. The work in J.P.F.A group also receives additional support from the Deutsche Forschungsgemeinschaft (DFG), FR 3746/3-1, FR 3746/5-1, FR 3746/6-1 and CRC205 (INST 269/886-1), the EU-H2020 (ERC-Consolidator, DeciFERR) and the Deutsche Jose Carreras Leukämie Stiftung (DJCLS 01 R/2022). M.F work was supported by the German Research Foundation (DFG, Deutsche Forschungsgemeinschaft) as part of the SFB1588 (grant ID 493872418 to M. Fischer), and the grants BA 6984/1-1 (to C. Bartenhagen), FI 1926/2-1 (to M. Fischer). The study was also supported by the Förderverein für krebskranke Kinder e.V. Köln (endowed chair to M. Fischer), Leverkusen hilft krebskranken Kindern e.V. (to M. Fischer), and the program “Netzwerke 2021”, an initiative of the Ministry of Culture and Science of the State of Northrhine Westphalia (NRW) for the CANTAR project (to M. Fischer).

S.M. was supported by Fundação de Amparo à Pesquisa do Estado de São Paulo [FAPESP, CEPID–Redoxoma 13/07937-8] and Conselho Nacional de Desenvolvimento Científico e Tecnológico [CNPq 313926/2021-2]. A.I was supported by Fundação de Amparo à Pesquisa do Estado de São Paulo [FAPESP, 2017/13804-1]. L.A.S. is supported by grants from the National Institute of Diabetes and Digestive and Kidney Diseases (#R01DK128390) and the Ingeborg v.F. McKee Fund from the Hawaii Community Foundation (#MedRes_2023_00002973). B.K.S. is supported by a grant from the National Institute of General Medical Sciences (#P20GM139753 – subproject 5203). U.S. is supported by Universitätsklinikum Bonn. F.M acknowledges the Fundação de Amparo à Pesquisa do Estado de São Paulo (FAPESP) projects, CEPID-Redoxoma 2013/07937-8, Young Investigator 2018/14898-2 and multi-user equipment 2015/10411-3. S.M. was supported by Fundação de Amparo à Pesquisa do Estado de São Paulo [FAPESP, CEPID– Redoxoma 13/07937-8] and Conselho Nacional de Desenvolvimento Científico e Tecnológico [CNPq 313926/2021-2]. A.M. is a MSCA-PF researcher funded by the EU (iroNKiller, GA: 101061974).

## Author contribution

Z.C performed CRISPR screen and most in vitro assays. CRISPR analysis was performed by U.Y. A.I contributed to in vitro studies and the characterization of PRDX6 enzymatic reactions and product analysis by MS with contributions from R.S.L, M.P.P, L.R.G and R.P.S. Those studies were supervised by L.E.N, F.M and S.M. K.K established and performed SELENOP uptakeassays and Duolink assays. T.C, contributed with the orthotopic xenografts with supervision from H.A. G.F established and performed the split GFP assays. T.N.X.S, F.P.F, M.F, contributed with performing and analysing in vitro assays. L.G.V, J.K.S and S.W contributed with computational analysis of SEPSH2/PRDX6 interacion and molecular dynamics simulation. M.F contributed with the studies using radiolabeled SELENOP, supervised by U.S. A.T, R.H, M.F, C.B, L.S, B.S contributed with reagentes and critical intelectual contribution. J.P.FA conceptualized the project. H.A, S.M and J.P.F.A supervised the study. Z.C, A.I and J.P.F.A wrote the first version of the manuscript. All authors read, contributed and agreed on the content.

## Materials and Methods

### Cell culture

The 4-hydroxytamoxifen (TAM)-inducible Gpx4−/− murine immortalized fibroblasts (Pfa1) (Kindly provided by Dr. Marcus Conrad), the human neuroblastoma cell line NMB and SK-N-DZ (purchased from ATCC), the human melanoma cell line A375 (purchased from ATCC), the human fibrosarcoma cell line HT1080 (purchased from ATCC), the human lung carcinoma cell line A549 (purchased from ATCC) and HT1975 (purchased from ATCC), the human embryonic kidney cell line HEK293T (purchased from ATCC) and the human hepatocellular carcinoma HepG2 (purchased from ATCC) were maintained in DMEM medium (Gibco) supplemented with 10% FBS (Gibco) and 100 U/ml penicillin/streptomycin (Gibco) at 37°C with 5% CO2. Cell lines were tested for mycoplasma contamination by the respective Multiplexion service (Heidelberg, Germany).

### Immunoblotting

Cell lysates were prepared with Radioimmunoprecipitation assay (RIPA) buffer containing protease inhibitor cocktail (Roche). Proteins were separated by SDS–polyacrylamide gel electrophoresis and electro-transferred onto polyinylidene difluoride membranes which then were subsequently blocked in 5% defatted milk in Tris-buffered saline-0.1 % Tween 20 (TBS- T) for 1 h at room temperature (RT). Primary antibodies were diluted in a solution of 5% bovine serum albumin (BSA) or 5% defatted milk in TBS-T. The membranes were incubated with primary antibody solution at 4°C overnight. Membranes were washed with TBS-T and incubated at RT for 2 h with horseradish peroxidase-labeled secondary antibody (1:3,000, anti-rabbit #7074, anti-mouse #7076 or anti-rat #7077, Cell Signaling) diluted in a solution of 5% milk in TBS-T. Proteins were detected using enhanced chemiluminescence substrate (107- 5061, BioRad). Images were acquired using detection systems Amersham ImageQuant 800 (Cytiva, EUA) or Azure 400 (Biozym, Germany).

Following antibodies were used against β-actin (1:5,000; A5441, Sigma-Aldrich), GPX4 (1:1,000; no. ab125066, abcam), LRP8 (1:1,000; ab108208, abcam), SCLY (1:50; IG-P1097, ImmunoGlobe GmbH), Flag (1:3,000; F3165, Sigma-Aldrich), PRDX6 (1:1000; GTX115262, GeneTex), MPST (1:1000; sc-376168, Santa Cruz Biotechnology), CBS (1:1000; sc-271886, Santa Cruz Biotechnology), CTH (1:1000, D4E9J, Cell Signaling) SELENON (1:1000; sc- 365824, Santa Cruz Biotechnology), VINCULIN (1:1000, sc-73614, Santa Cruz Biotechnology), DSRed (1:1000, 632496, TaKaRa).

### Molecular cloning

The gene fragments of interest were synthesized from Genescript and cloned into lentiviral vector p442-IRES (kindly provided by Dr. Marcus Conrad) for gene expressing via restriction digestion and Clone Express MultiS One Step Cloning Kit (Vazyme).

Individual sgRNAs were cloned into the pLenti-CRISPR-V2 (Addgene #98293) lentivectors for CRISPR Knock-out (CRISPR-KO) experiments via restriction digest of the respective lentivector with BsmBI (NEB, Cat. No. R0739) and T4 ligase (NEB, Cat. No. M020L). Oligonucleotides (Eurofins Genomics) with the sgRNA sequences and complementary overhangs were phosphorylated, annealed, and inserted into the respective lentiviral delivery vector (Table 1).

**Table 1.**
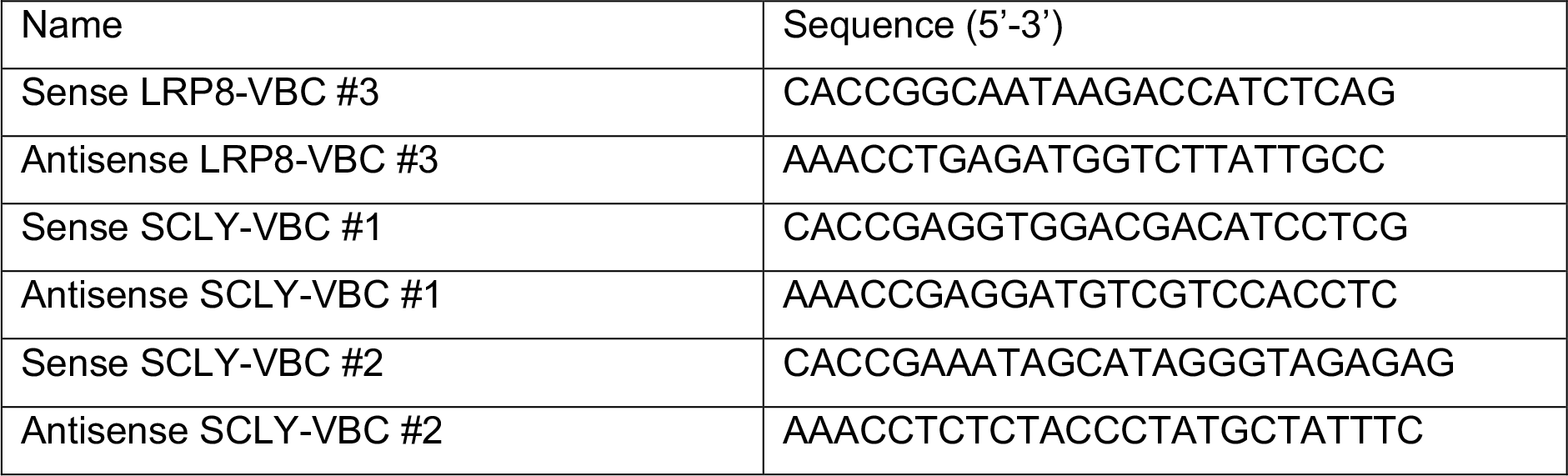
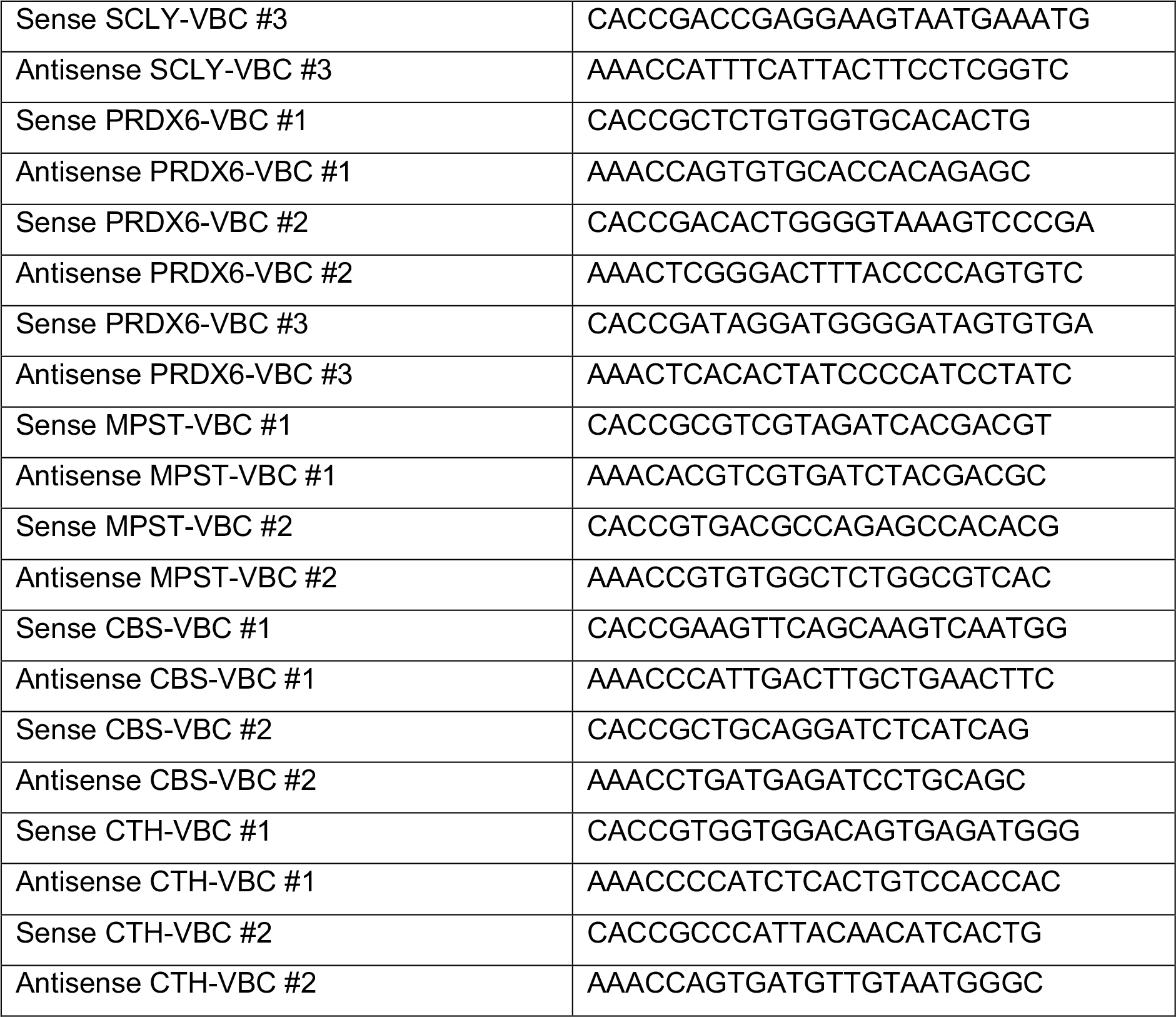
Oligonucleotides used for cloning of individual gRNAs.

### Lentivirus production

Lentivirus was produced with a third-generation packaging system and X-tremeGENE HP DNA Transfection Reagent (Sigma-Aldrich, 6366546001). HEK293T cells were transfected with the lentiviral transfer plasmids, two packaging plasmids (pMDLg/pRRE, Addgene #12251 and pRSV-Rev, Addgene #12253) and the envelope expressing plasmid (pCMV-VSV-G, Addgene #8454). Virial supernatant was harvested 48 h post-transfection, snap frozen with liquid N2 and stored at −80°C. Alternatively, the following constructs: p442-IRES-Mock, p442- IRES-LRP8-Flag, p442-IRES-FSP1, p442-IRES-SCLY, p442-IRES-PRDX6, p442-IRES- PRDX6 (C47S), p442-IRES-PRDX6 (S39A) and p442-IRES-PRDX6 (S39Y) were used for the generation of the cell expressing. All experimental procedures for lentivirus production and transduction were performed in a biosafety level 2 laboratory.

### Cell line generation from genome-wide CRISPR screen

Polyclonal SK-N-DZ cells were transiently transfected with the plasmid pLenti-CRISPR-V2- SCLY sgRNA #3. After 3 days selection with puromycin (1 μg/ml), cells were grown by limiting dilution. Clonal cell lines were established and tested by Western blot. Two clonal cell lines were selected for further experiments. To confirm, there was no sgRNA integration into the genome. The puromycin treatment was applied again to the selected clonal cell lines.

### Genome-wide CRISPR screen to identify potential enzyme in Selenocysteine metabolism

The monoclonal SK-N-DZ SCLY^KO^ cell line was transduced with Toronto KnockOut (TKO) CRISPR Library - Version 3 (Addgene, 90294) at MOI of 0.3. A total of 38 million cells were transduced to achieve > 500 cells per sgRNA. After selection with 1 μg/ml puromycin and recovery, the SCLY-KO cell line was splited into two conditions and maintained at the sgRNA coverage of > 500. Cell pellets were harvested after 2 weeks of passaging. Genomic DNA purification and Sequencing. The mapping of raw sequencing reads to the reference library, and computing of enrichment scores and P-values were processed using the MaGeCK pipeline. The MaGeCKFlute was used for the identification of gene hits and associated pathways(Yang et al., 2020).

### Generation of HepG2 conditioned medium and SELENOP enrichment

HepG2 cells were seeded to a 15-cm^2^-round dish with standard DMEM medium (10% FBS+ 1% Pen/Strep). The medium was changed to FBS-free medium supplemented with 50 or 200 nM Na2SeO3 when HepG2 cells achieved ∼70–80% confluent. After 48 h incubation, the medium was collected and briefly spined down to clarify the supernatant. The supernatant was transferred to the centrifugal filter (Amicon Ultra-15, 30K. UFC903024) and centrifuged for 10 min at 3,000 g. After the supernatant was centrifuged through the centrifugal filter, 2 ml of FBS-free medium was used to resuspend the concentrated fraction from the filter. The target cell lines were plated and treated with HepG2 concentrated medium in 6-well plates in the absence of FBS. After 24-h incubation, the cell lysates were collected.

### ^75^Se direct labelling and Hot ^75^Se-SELENOP labelling

For Na2^75^SeO3 direct labelling, SK-N-DZ LRP8^OE^ with PRDX6^WT^/SCLY^WT^, PRDX6^KO^/SCLY^WT^, PRDX6^WT^/SCLY^KO^ and PRDX6^KO^/SCLY^KO^ cell lines were plated in 6 cm-dishes in FBS-free medium (DMEM) + 500 nM Liproxstatin-1 supplemented with 6 μCi of Na2^75^SeO3 (Polatom). After 48 h incubation, cells were collected in RIPA buffer and separated by 12% SDS– polyacrylamide gel electrophoresis. Screens were exposed for 1 week.

HepG2 cells were seeded in 4 T75 flasks in FBS-free medium (DMEM-F12) supplemented with 15 μCi of Na2^75^SeO3 (Polatom) per flask. After 48 h incubation, the media was collected and ^75^Se-SELENOP was concentrated by centrifugation in Amicon tubes (cut off 10K) followed by desalting in Sephadex G25 columns. For ^75^Se-SELENOP labelling, the target cell lines were plated in 6 cm-dishes in FBS-free medium (DMEM) + 500 nM Liproxstatin-1 supplemented with ^75^Se-SELENOP. After 48h of cultivation, cells were collected in RIPA buffer and loaded on 12% SDS- polyacrylamide gels. Screens were exposed for 1 week.

### Viability assay

The impact of compounds or gene deficiency on cell viability was assayed with the Alamar blue assay. To determine the cell viability, 3,000 cells were seeded in 96-well plate with standard medium (10% FBS + 1% penicillin/streptomycin) 4 h ahead the treatment. After 72 h incubation, cell viability was analyzed using the Alamar blue assay following the manufacturer’s instructions. Alamar blue solution was made by dissolving of 1 g resazurin sodium salt (Sigma) in 100 mL sterile PBS and sterile filtrated through a 0.22 µm filter. After 2-4 h incubation, viability was assessed by measuring the fluorescence using a 540/35 excitation filter and a 590/20 emission on a Spark® microplate reader (Tecan, Zürich, Switzerland). For clonogenic assay, 500 cells were plated in 6-well plate for 14 days under respective conditions. Then, the colonies were stained with 0.01% (w/v) crystal violet-PBS solution.

### Assessment of lipid peroxidation using BODIPY C-11 (581/591)

100,000 cells per well were seeded in 6-well plate, one day prior to the experiment. Subsequently, the cells were incubated with 1 µM BODIPY C-11. After 30 min incubation at 37 °C inside the tissue culture incubator, the cells were harvested and analyzed with FACS Canto II cell sorter.

### Expression, purification and quantification of Prdx6 from *Rattus norvegicus*

N-terminal, His-tagged PRDX6 from rat (gene cloned into pET15b NdeI and XhoI restriction sites) as produced in the *Escherichia coli* strain Origami *(DE3) (*Novagen). 20 µL of Prdx6- Pet15b transformed Origami (*DE3*) cells were inoculated in 50 mL of lysogeny broth (LB) medium. To the same medium, we added kanamycin (15 µg/mL), tetracycline (12.5 µg/mL) and ampicillin (50 µg/mL). Bacteria were grown overnight at 37 °C and 200 rpm using an orbital shaker. Bacteria culture was then diluted to OD600 = 0.2 in 500 mL of LB medium containing 15 µg/mL kanamycin, 6.25 µg/mL tetracycline and 100 µg/mL ampicillin. This culture was grown to OD600nm = 0.8 at 37 °C and 200 rpm. IPTG (500 µM) was then added, and the culture was kept under agitation at 20 °C overnight. The resulting pellet was washed with Milli-Q^®^ water and centrifuged again (10 min, 4 °C, 5000 rpm), and finally stored in a - 20 °C freezer.

For protein extract preparation, bacterial pellet was resuspended in 50 mM sodium phosphate buffer (pH 7.4) with 500 mM NaCl, using 20 mL of buffer for each 500 mL of growth medium. 50 µL of lysozyme (10 mg/mL) and 500 µL of protease inhibitor cocktail (SIGMAFAST™ Protease Inhibitor Cocktail Tablets, EDTA-free) were added to the solution, which was stirred for 10 min at room temperature. Bacteria were subsequently sonicated for 5 minutes (amplitude 30%, pulse on for 20 seconds, pulse off for 60 seconds). After cell lysis, streptomycin sulfate (1%) was added for nucleic acid precipitation for 20 min with agitation. Cells were then centrifuged (15000 rpm) for another 30 min at 4 °C . The precipitate was discarded and the supernatant was filtered through a 0.45 µm membrane (Millipore/Merck KGaA, USA).

Recombinant PRDX6 was purified by nickel affinity chromatography, using a 5 mL column (Ni-NTA Superflow Cartridges, QIAGEN, Netherlands) with an imidazole gradient: Buffer A (20 mM sodium phosphate buffer, pH 7.4, with 500 mM NaCl) and Buffer B (20 mM sodium phosphate buffer, pH 7.4, with 500 mM NaCl and 500 mM imidazole). This procedure was performed in an ÄKTA FPLC System (Amersham Pharmacia Biotech, United Kingdom). Fractions were collected and evaluated on a 14% SDS-Page (Molecular weight of recombinant His tagged PRDX6 is 26.850 Da). Selected fractions were pooled and concentrated to a final 1.5 mL volume by centrifugation (4900 rpm, 30 min, 4 °C) using Amicon^®^ Ultra-15 10 KDa device (MilliporeSigma, USA).

Excess imidazol was removed by gel filtration_through a HiTrap® Desalting column (GE Healthcare, USA). Then, PRDX6 concentration was determined spectrophotometrically (Ɛ280nm = 2.556×10^4^ M^−1^ cm^−1^).

### Identifying rPrdx6 modifications by selenium compounds

#### rPrdx6 reduction with dithiothreitol (DTT)

Both wild-type (WT) and mutant (C47S) Prdx6 from *Rattus norvegicus* (rPrdx6) were initially reduced with DTT (three-fold excess, approximately 1 mM) for 2 h at 37 °C in a water bath. After reduction, excess DTT was removed using ‘Amicon® Ultra 0.5 mL 10 K centrifugal filters’ and Milli-Q^®^ water; six centrifugation steps were performed at 10000 g for 5 minutes. Final protein concentration was then estimated by measuring the UV absorbance at 280 nm using a SpectraMax^®^ M2 Reader (Molecular Devices, USA).

#### Incubation of rPrdx6 with selenium and sulfur compounds

30-40 µM of WT or C47S rPrdx6 were incubated with 1 mM sodium selenide (Na2Se), 800 µM sodium selenite (Na2SeO3), 800 µM L-selenocystine, 800 µM L-selenomethionine, and 800 µM or 3.2 mM L-cystine at 37 °C and 500 rpm using an Eppendorf ThermoMixer^®^ (Eppendorf, Germany). Incubation times varied according to the reagent. Incubations had a final volume of 200 µL and were performed at both acidic or neutral pH (5.0 or 7.4). We used 100 mM sodium phosphate buffer for both pH values (5.0 and 7.4); buffers were pre-treated with 10 µg/mL catalase for 2 hours for H2O2 removing. We also performed co-incubations of selenium compounds with (two or four-fold excess) reduced L-glutathione (GSH). L-selenocystine and L-selenomethionine were acquired from Cayman Chemicals (USA); all the others were provided by Sigma-Aldrich (USA).

#### Non-digested protein analysis through mass spectrometry

For mass spectrometry analysis, excess of buffer and selenium/sulfur reagents was removed from the protein reaction products using ‘Amicon® Ultra 0.5 mL 10 K centrifugal filters’ and Milli-Q^®^ water; two centrifugation steps were performed at 10000 g for 5 minutes. ‘Washed’ samples were diluted to approximately 30-40 µM and then injected into the Mass Spectrometer (SCIEX TripleTOF 6600) by direct infusion (10 µL/min) together with a flow rate of 0.2 mL/min of water and acetonitrile (1:1) and 0.1% formic acid. Parameters in Table 2 were used for data acquisition (software: Analyst^®^TF 1.7.1) and protein reconstruction/deconvolution (software: PeakView^®^ 2.2).

**Table 2.**
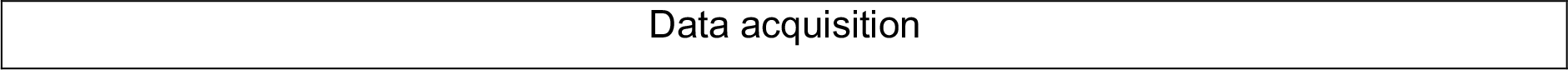
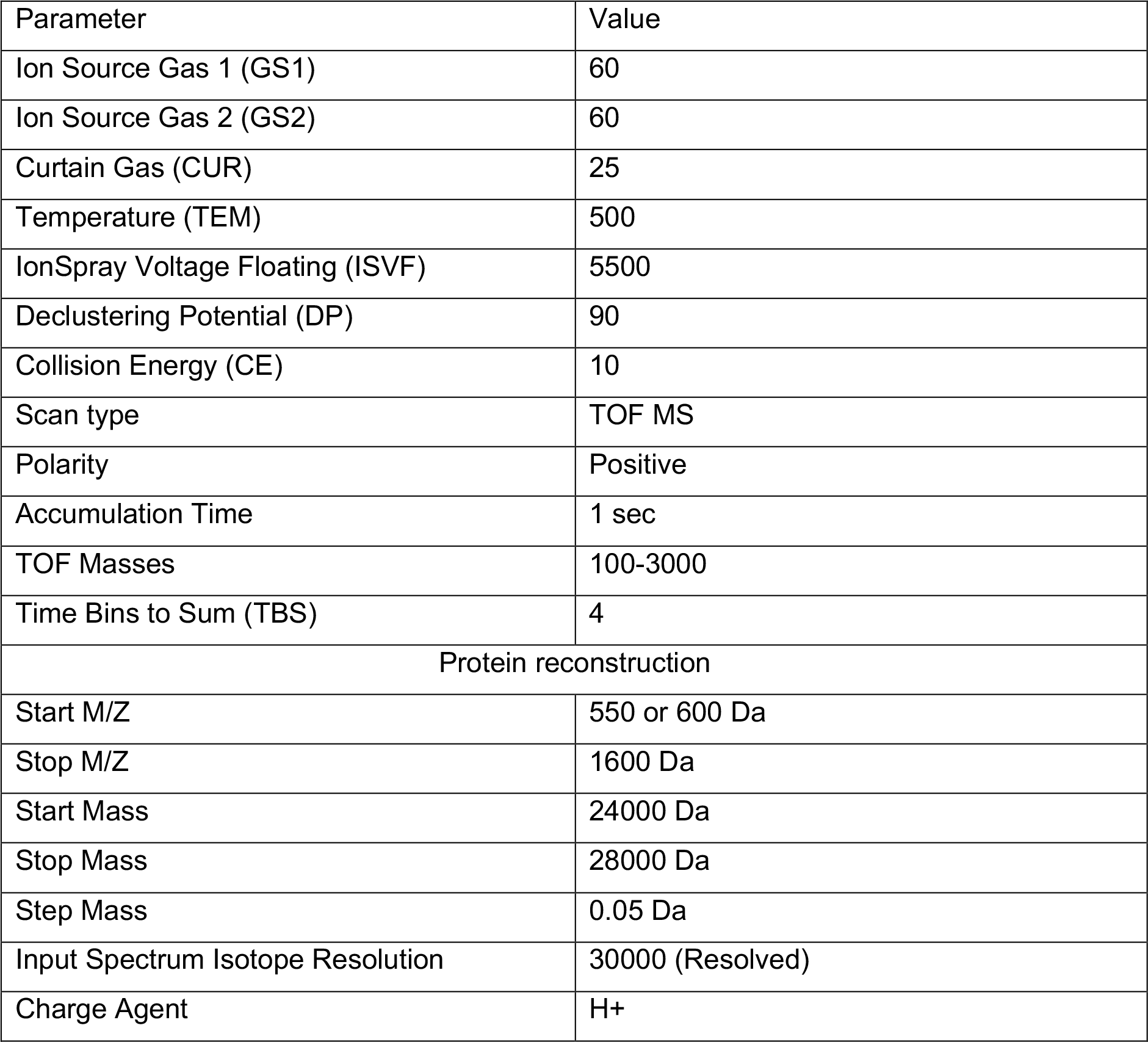
Parameters for mass spectrometry analysis of non-digested rPrdx6 and protein reconstruction/deconvolution.

| Parameter | Value |
| --- | --- |
| Ion Source Gas 1 (GS1) | 60 |
| Ion Source Gas 2 (GS2) | 60 |
| Curtain Gas (CUR) | 25 |
| Temperature (TEM) | 500 |
| IonSpray Voltage Floating (ISVF) | 5500 |
| Declustering Potential (DP) | 90 |
| Collision Energy (CE) | 10 |
| Scan type | TOF MS |
| Polarity | Positive |
| Accumulation Time | 1 sec |
| TOF Masses | 100-3000 |
| Time Bins to Sum (TBS) | 4 |
| Protein reconstruction |  |
| Start M/Z | 550 or 600 Da |
| Stop M/Z | 1600 Da |
| Start Mass | 24000 Da |
| Stop Mass | 28000 Da |
| Step Mass | 0.05 Da |
| Input Spectrum Isotope Resolution | 30000 (Resolved) |
| Charge Agent | H+ |

### Determining the oxidation kinetics (peroxidase activity) of rPrdx6

#### Photooxidation of phosphatidylcholine (PC)

Photooxidation was carried in a double neck glass balloon containing 70 mg of EggPC (Sigma P3565) diluted on 7 ml of chloroform. 200 ul of a 2.5 mg/ml solution of methylene blue was used as a sensitizer. Light source was a 500 W tungsten lamp. Reaction mixture was kept on ice, oxygen rich atmosphere, and constant agitation for 4 h. Progress of the reaction was followed by RP-TLC C18, mobile phase chloroform:methanol:water (5:10:0.5 v/v/v). Iodine vapor was used as visualization reagent (Alegria et al., 2017).

#### Isolation of mono-hydroperoxide PC (PC-OOH)

Mono-hydroperoxide PC (PC-OOH) was isolated by high performance liquid chromatography (HPLC Prominence, Shimadzu, Kyoto, Japan) using a semi-preparative column (Phenomenex, Luna C18(2), 5µm, 100 Å, 250 x 10 cm). Mobile phase consisted of water for phase A, and methanol for phase B, in a linear gradient as follows: 97% B (0 to 8 min); 97 to 100% (8 to 12 min); 100% to 97% (12 to 14 min); 97% B (14 to 15 min). The oven temperature was 32 °C, and the flow rate was set to 4,7 mL.min-1. The fractions containing PC-OOH were collected and the solvent evaporated. PC-OOH was diluted in ethanol and stored at -80 °C. Quantification of hydroperoxides was performed using the iodometric assay. Oxidation of PRDX6 can be measured by changes in the intrinsic fluorescence of the protein when it is incubated with peroxides, which is monitored using stopped-flow spectroscopy.

#### Determining the oxidation kinetics of rPrdx6

An aliquot of 335 µM recombinant PRDX6 in 20 mM sodium phosphate buffer with 100 mM NaCl was reduced with approximately 1 mM DTT for 2 h at 37 °C (water-bath), in the dark. After reduction, excess DTT was removed through six washes with Milli-Q^®^ water using an Amicon^®^ Ultra 0.5 mL 10 K centrifugal filter (centrifugation: 10000 g, 5 minutes, 5 °C). Re-quantification of the protein was performed in a quartz cuvette at 280 nm using a SpectraMax^®^ M2 Reader (Molecular Devices, USA). To calculate the protein concentration, its molar extinction coefficient was determined as 22460 M^-1^cm^-1^ by the ProtParam tool of the ExPASy Proteomics Server (www.expasy.org). An aliquot of the protein was incubated with 4,4′- dithiodipyridine (Aldrithiol™-4) (3-fold excess) for thiol quantification at 324 nm in the presence of 0.15% SDS. A molar extinction coefficient of 21400 M^-1^cm^-1^ was used in the calculations for 4-thiopyridone – the reduced form of Aldrithiol™.

We used hydrogen peroxide (H2O2), linoleic acid (mono)hydroperoxide (LA(OOH)1) and PC- OOH as substrates for PRDX6’s peroxidase activity. The concentration of H2O2 stock solution was determined at 240 nm in an UV-Vis spectrophotometer (Cary 60, Agilent Technologies, CA, USA), considering a molar extinction coefficient of 43.6 M^−1^cm^−1^. Dilutions of the protein and the peroxides were prepared in 100 mM sodium phosphate buffer (pH 5.0 or 7.4) containing 100 µM diethylenetriaminepentaacetic acid (DTPA). Buffer solutions were pre-treated with 10 µg/mL catalase for 2 h to remove any trace of H2O2, and then filtered by centrifugation (2000 g, 15 min, 5 °C) using Amicon® Ultra-15 Centrifugal Filters (30 kDa cutoff). PRDX6 was used at a final concentration of 0.5 µM and mixed with increasing concentrations of H2O2 (0.5 to 2.5 µM) and lipid peroxides (15 to 150 µM) in a stopped-flow instrument (Applied Photophysics SX20MV, United Kingdom), with excitation (λ) at 280 nm and temperature of 25 °C. A final percentage of 5% ethanol was necessary to solubilize lipid peroxides in buffer solutions.

Observed rate constants (*kobs*) for rPrdx6’s fluorescence change was determined by fitting data to single exponential equations. Second order rate constants were obtained by the slope in the linear fitting of the *kobs* versus peroxide concentration.

### Homology modeling and rat Prdx6 structure preparation

The structure of Rattus norvegicus Prdx6 (rat Prdx6) was modelled using the homology modelling package SWISS-MODEL (Waterhouse et al., 2018). The crystal structure of the human Prdx6 (91.5% identity with rat Prdx6; PDB code: 5B6M; resolution: 2.50 Å) (Kim et al., 2016) was used as a template to build the model. The quality of the generated model was assessed using the quality assessment tools of SWISS-MODEL, including: (i) analysis of Ramachandran plots to visualize energetically favoured regions for backbone dihedral angles of amino acid residues of the protein model and (ii) analysis of QMEAN score (Benkert et al., 2009), which is based on geometrical properties and provides both global and local quality estimates.

### Molecular dynamics (MD) simulations

The rat Prdx6 structure was obtained by homology modeling, as described above. All protein hydrogen atoms were added to each structure using the PDB2PQR tool (Jurrus et al., 2018), with AMBER forcefield. The predicted predominant protonation states of ionizable groups from titratable residues were assigned at pH = 7.0 or at pH = 5.0, using PROPKA(Olsson et al., 2011; Sondergaard et al., 2011) for prediction of pKa values. All MD simulations were carried out using AMBER v.18 package (Salomon-Ferrer et al., 2013). The topology and coordinate files were built using the LEaP module of AmberTools (Salomon-Ferrer et al., 2013), with AMBER ff14SB forcefield (Maier et al., 2015) parameters. The system was solvated using a TIP3P explicit water solvent model (Jorgensen et al., 1983), with a total charge of zero, in a cubic box with at least 10 Å between the protein surface and the box border.Energy minimization was performed with steepest descent method and conjugate gradient in four steps, gradually reducing the restraints on the protein atoms to their initial positions. In the three first steps, the restraint force constants were defined as 500, 100, and 5 kcal.mol-1.Å-², respectively. In the last step, all restraints were removed, and the entire system was minimized. Each minimization step was carried out for a maximum of 1500 iterations, with 500 iterations of steepest descent minimization followed by 1000 iterations of conjugate gradient minimization.Next, the system was allowed to heat up from 10 to 300 K, through 1 ns of MD simulation, using 2 fs time steps. A Langevin thermostat with a collision frequency of 1.0 ps-1 and constant volume periodic boundaries were used. Protein atoms were restrained to their initial positions, with a restraint force constant of 50 kcal/(mol.Å²). Afterward, the system was equilibrated at 300 K, through 20 ns of MD simulation, using a Langevin thermostat with a collision frequency of 1.0 ps-1 and constant pressure of 1 bar, with restraints on the protein atoms positions gradually released during four steps (restraint force constants of 50, 10, 2 and 0 kcal.mol-1.Å-², respectively). The system was treated under periodic boundary conditions. Particle-mesh Ewald (PME) electrostatics and a cut-off of 10 Å for non-bonded interactions were used. A time step of 2 fs was applied for integration and coordinates were written every 2 ps. Finally, production MD simulation runs (100 ns) were carried out, using the same setup as described for the last equilibration step. For each system, three independent simulations were performed, starting from different initial velocities distributions. The MD trajectories were analyzed using the cpptraj module of AmberTools v.18 and/or the MD trajectory analysis tools of VMD v.1.9.2 (Humphrey et al., 1996). Solvent-accessible surface areas (SASA) for the protein’s active site and for Cys-47 were calculated using the “linear combinations of pairwise overlaps” (LPCO) method (Weiser et al., 1999) implemented in cpptraj. GraphPad Prism v.9 was used for plotting. Molecular visualization was performed with PyMOL v.2.5.0 (https://pymol.org/2/) or VMD v.1.9.2 (https://www.ks.uiuc.edu/Research/vmd/) (Humphrey et al., 1996).

### Split SF-Ch assay

The constructs for SF-Ch2 assays were obtain from Addgene (plasmid #82604 and #82602). Constructs were then cloned and expressed using the lentiviral system as described above. Confocal images were obtained using a Leica SP8 microscope with a HCX PL APO CS 63x 1.4 NA oil objective.

### Duolink proximity ligation assay

The interaction between PRDX6 and SEPHS2 in SK-N-DZ WT and SK-N-DZ *PRDX6 ^KO^* cells was observed using Duolink in situ proximity ligation assay (PLA) kit (DUO92101, Sigma-Aldrich) in the presence or absence of 600nm Selenocysteine. The cells were fixed in 4% PFA for 10 min at room temperature and permeabilized with 0.01% Triton X, before being blocked with a blocking solution. The cells were incubated with primary antibodies targeting for respective proteins for 1 h at 37 °C, followed by incubation with PLA probes for 1 h at 37 °C in a humidified chamber. After three washes, a ligation-ligase solution was added and incubated for 30 min at 37 °C. The slides were incubated for 100 min in an amplified polymerase solution at 37 °C in the dark. Finally, the cells were stained with a mounting medium containing DAPI.

Fluorescence microscope was used to capture the fluorescence images (Cell Discoverer, Germany).

### In vivo orthotopic mouse experiments

Following German legal regulations, all studies involving mice and experimental protocols were approved by the Regierungspraesidium Karlsruhe, the state’s governmental review board, under the authorization number G-176/19. The studies were carried out in accordance with the guidelines of the German Cancer Centre Institute. NXG mouse strains were used in the study. Experiments were conducted using female mice aged 3-4 months. Mice were kept in separately ventilated cages under temperature and humidity control. The cages were provided with an improved habitat including bedding material. To generate orthotopic mouse models for neuroblastoma, 2 × 10^5^ SK-N-DZ with *PRDX6^WT^/SCLY^WT^*and *PRDX6^KO^/ SCLY^KO^* cells were transplanted into the right adrenal gland after the surgical site was prepared. Cells were resuspended in a 1:1 (vol/vol) mix of growth factor-reduced matrigel (Corning) and PBS. A total of 20 μl of this cell suspension was injected into the right adrenal gland of anaesthetized mice. Following tumour cell transplantation, we used an IVIS Spectrum Xenogen device (Calliper Life Sciences) to monitor the mice for bioluminescent signals indicating tumour growth. One week after injecting the cells, we noticed a clear signal from the tumours. Animal health was evaluated on a regular basis, and the mice were terminated as soon as they met the abortion criteria outlined in the procedure. The sample size was calculated with the help of a biostatistician using R version 3.4.0. Power analysis assumptions were as follows: α error = 5%, β error = 20%. Mice were randomly assigned to treatment groups prior to treatment. If animals had to be sacrificed before reaching the predetermined endpoint (due to weight loss or other termination circumstances), they were eliminated from further studies. All animal experiments were blinded during experiments and follow-up assessments.

### Prediction of PRDX6-SEPHS2 binding interface

Computational structural modeling was used to predict the putative binding interface between PRDX6 and SEPHS2. The experimentally determined structures for PRDX6 (PDB ID 5B6M) and with a sulfuric acid state (PDB ID 5B6N), which omitted the C47 active site and the AlphaFold prediction of a truncated version of SEPHS2, which omitted the U60 active site (B4E093) and a full-length version based on the sequence (Uniprot ID Q99611) were subjected to complex prediction using ClusPro(Desta et al., 2020). Active sites for were defined as R132, H39 and C471 for PRDX6, and U60 and G319-H3252 for SEPHS2. Every pairing possible of the two versions of PRDX6, and of SEPHS2 were subjected to ClusPro. ClusPro is unable to run with Q99611 due to the presence of U60, so the U60 was manually replaced by C60. ClusPro resulted in up to 29 different poses and the distance between active sites was found across all poses for every combination. For combinations that included PRDX6 sulfuric acid state, which would not contain the C47 active site, the peptide bond between H39 and P40 was used to measure from, and for the combinations that included the shorter form of SEPHS2, which would not have contained the C60 active site, the peptide bond between G319 and F320 was used to measure from. Distances were measured and structures visualized using PyMol software (https://pymol.org/).

---

**Figure S1.**
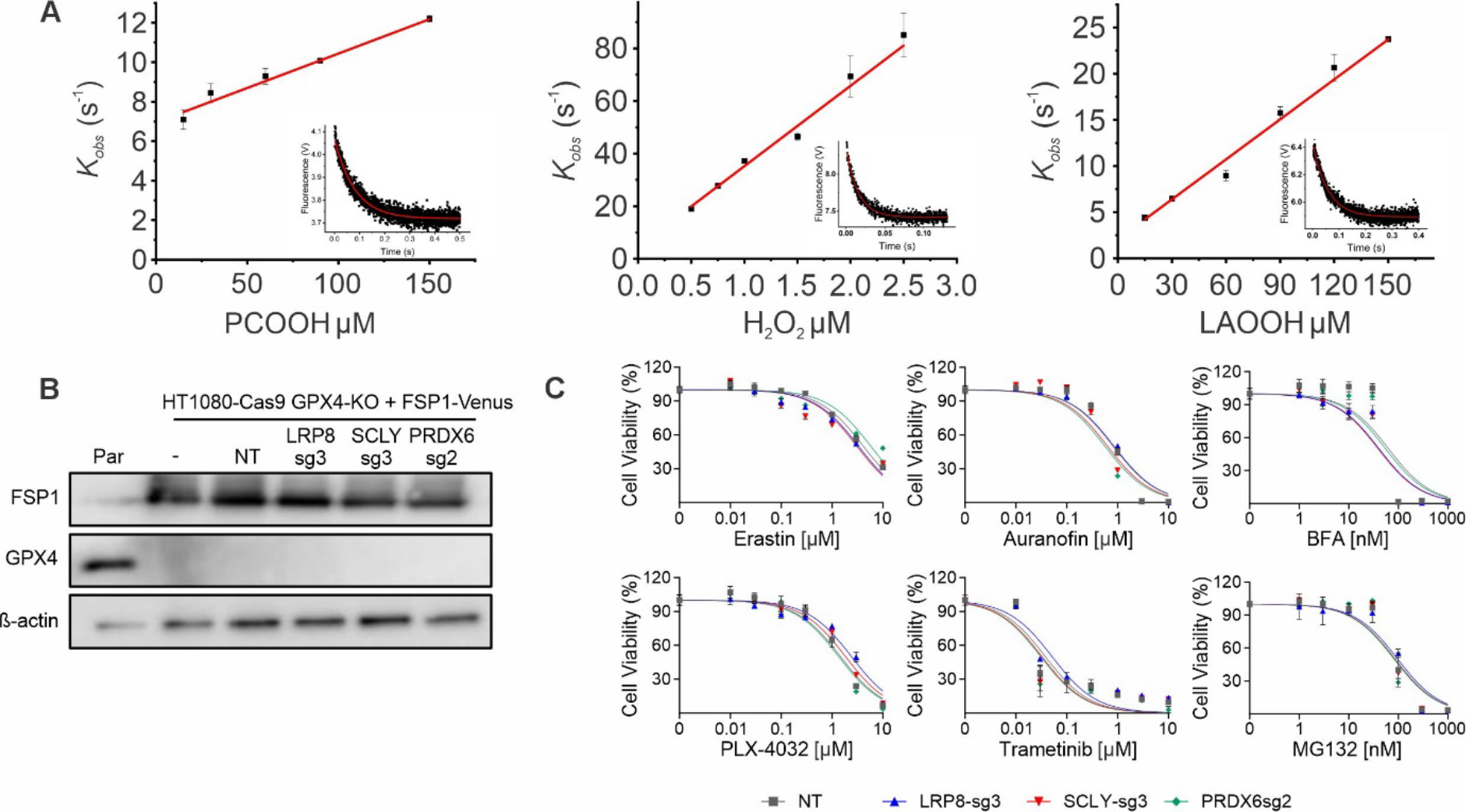
A. Monitoring of PRDX6 reaction with the indicated substrates over time by the variation of the intrinsic protein fluorescence. The *kobs* of the first rapid fluorescence decay were plotted against substrate concentration and the second-order rate constants calculated from the slope at pH 7.4; Representative exponential fittings are shown in the insets with 150 μM PCOOH, 2 μM H2O2, or 90 μM LAOOH. B. Immunoblot analysis of FSP1 and GPX4 in the wild-type, HT1080 with *GPX4* deficient and FPS1-Venus overexpression transduced with sgRNAs targeting *LRP8*, *SCLY* and *PRDX6*. C. The viability of HT1080 with *GPX4* deficient and FPS1-Venus overexpression with sgRNAs targeting *LRP8*, *SCLY* and *PRDX6* in the present of increasing concentration of Erastin, Auranofin, BFA, PLX-4032, Trametinib and MG132. Data are the mean ± SD of n=3 wells of 96-well plate.

**Figure S2.**
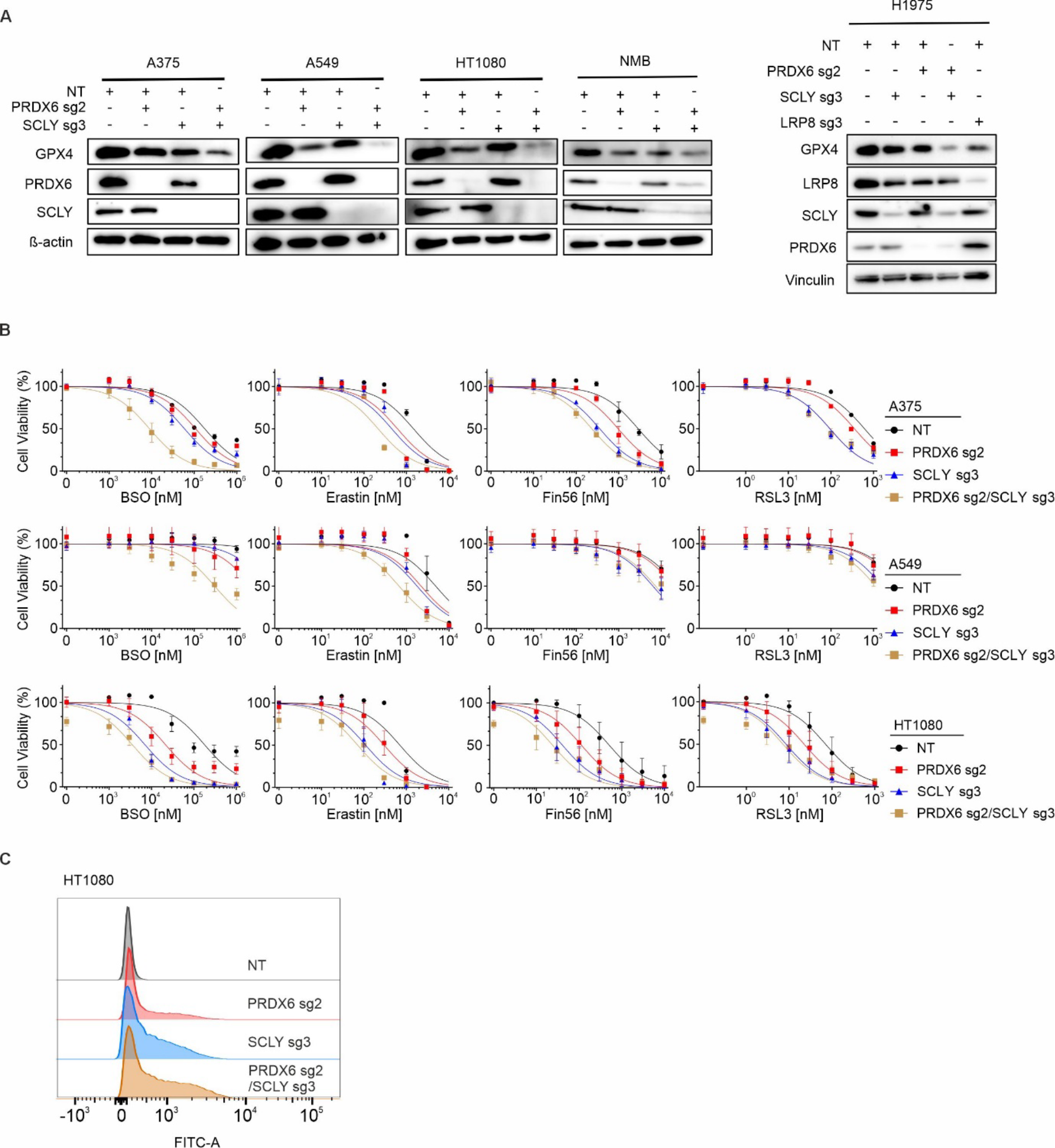
A. Immunoblot analysis of SCLY, PRDX6 and GPX4 in a panel of cell lines (A375, A549, HT1080, NMB and H1975) transduced with sgRNA targeting *PRDX6* and *SCLY*. B. Viability of A375, A549 and HT1080 transduced with sgRNAs targeting *SCLY* and *PRDX6* in the present of increasing concentration of RSL3, Fin56, Erastin and BSO. Data are the mean ± SEM of n=3 wells of 96-well plate from three independent experiments. C. Flow cytometry analysis of BODIPY 581/591 C11 oxidation in HT1080 transduced with sgRNA against *PRDX6* and *SCLY*.

**Figure S3.**
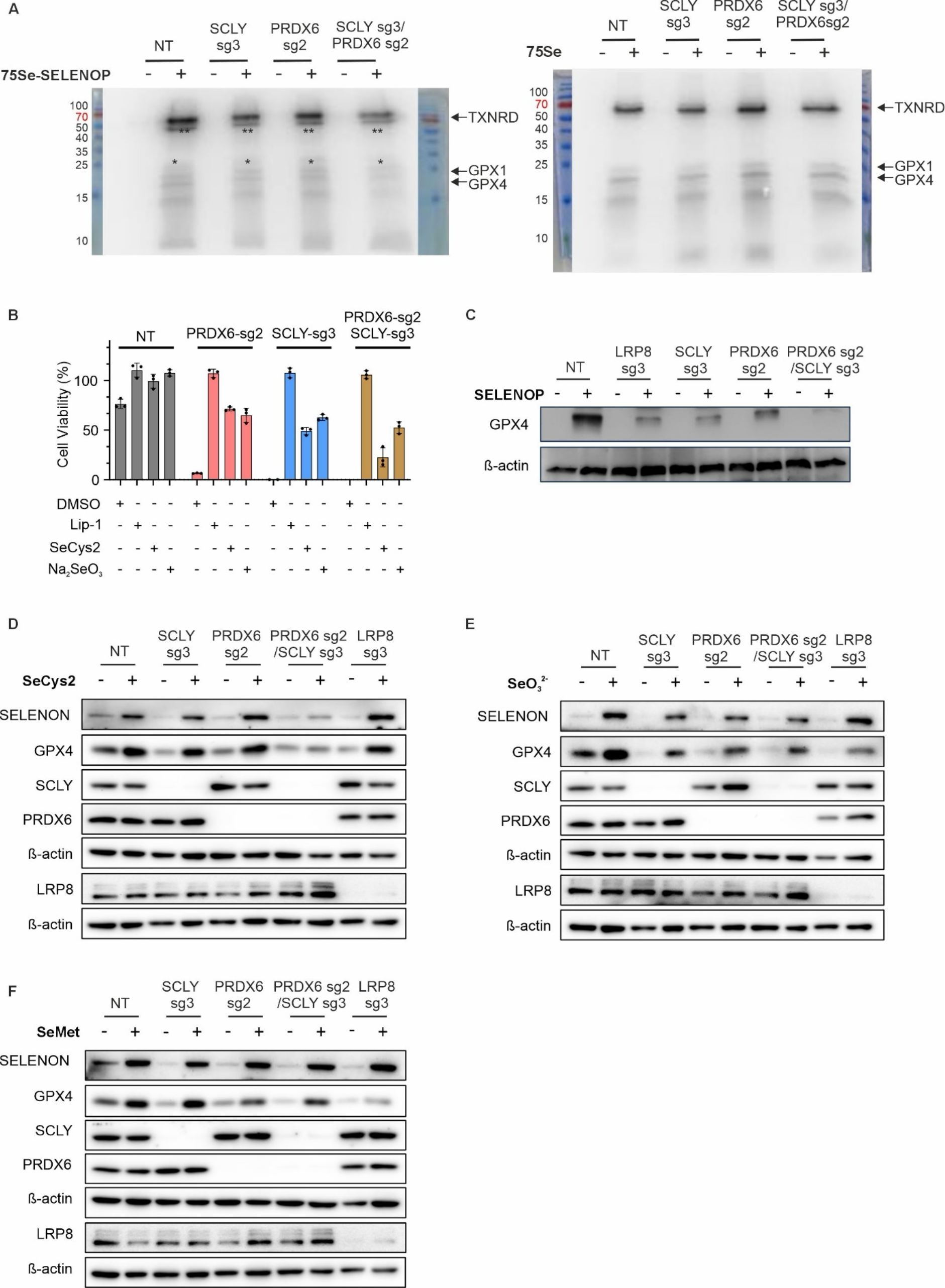
A. Autoradiography of ^75^Se metabolically labelled proteins. Left panel, global ^75^Se- selenium incorporation from ^75^Se-SELENOP into SK-N-DZ overexpressing LRP8-Flag (LRP8^oE)^ and deficient in *PRDX6*, *SCLY,* and *PRDX6/SCLY*. Right panel, global ^75^Se-selenium incorporation from Na2Se^75^O3 into SK-N-DZ LRP8^oE^ cell line and deficient in *PRDX6*, *SCLY,* and *PRDX6/SCLY* deficient with (^75^Se). ** ^75^Se-SELENOP and * fragment of ^75^Se-SELENOP. B. Viability of SK-N-DZ transduced with sgRNA against PRDX6, SCLY and both in the present of 500 nM Lip1, 500 nM L-selenocystine and 500 nM Na2SeO3. C. Immunoblot analysis of GPX4 in the wild-type and NMB transduced sgRNA against *PRDX6, SCLY* and *LRP8* with supplement of SELENOP. D. Immunoblot analysis of GPX4, SELENON, PRDX6, SCLY and LRP8 in the NMB transduced with sgRNAs targeting *LRP8*, *SCLY* and *PRDX6* with or without supplement of 50 nM L-selenocystine. E. Immunoblot analysis of GPX4, SELENON, PRDX6, SCLY and LRP8 in the NMB transduced with sgRNAs targeting *LRP8*, *SCLY* and *PRDX6* with or without supplement of 50 nM Na2SeO3. F. Immunoblot analysis of GPX4, SELENON, PRDX6, SCLY and LRP8 in the NMB transduced with sgRNAs targeting *LRP8*, *SCLY* and *PRDX6* with or without supplement of 5 µM L-selenomethione.

**Figure S4.**
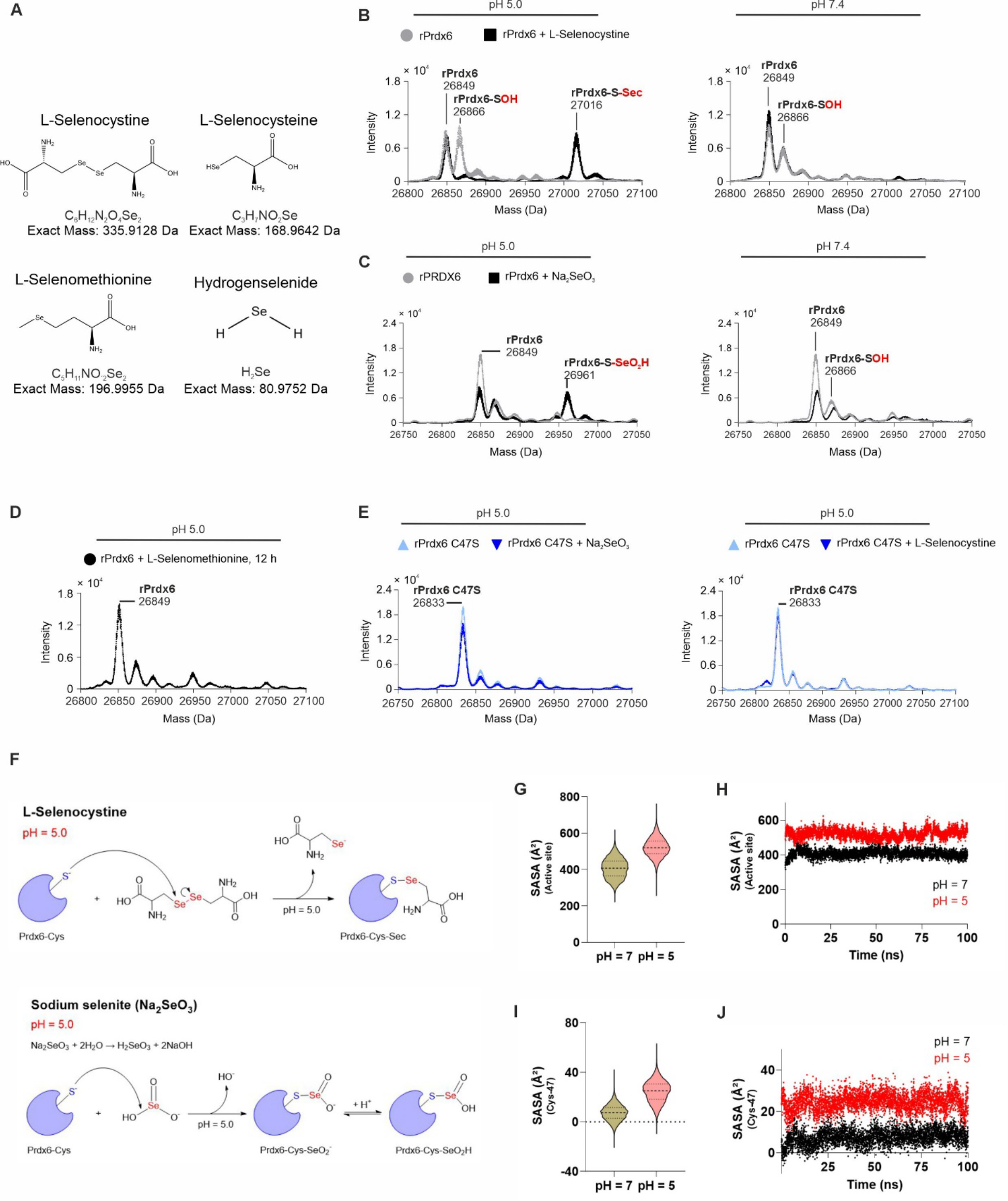
A. Molecular formulas and exact mass values obtained from ChemDraw (PerkinElmer, USA) for L-selenocystine and L-selenocysteine, L-selenomethionine; and Hydrogen Selenide. B. Incubation of 40 µM rPrdx6 with 800 µM L-Selenocystine in 100 mM PBS buffer (pH 5.0 and pH 7.4, 500 rpm, 37 °C). C. Incubation of 30 µM rPrdx6 with 300 µM Na2SeO3 for 30 minutes (pH 5.0 and pH 7.4, 500 rpm, 37 °C). D. Incubation of 40 µM rPrdx6 with 800 µM L-selenomethionine (pH 5.0, 500 rpm. 37 °C) E. Incubation of 40 µM rPrdx6 with 800 µM Na2SeO3 or 800 µM L-selenocystine in 100 mM PBS buffer (pH 5.0 and pH 7.4, 500 rpm, 37 °C). F. Schematic representation of rPrdx6 and L-selenocystine or Na2SeO3. G. Violin plots showing the distributions of the rPrdx6 active site’s SASA values among the MD simulations at pH = 7 and at pH = 5. Data represents the values from three independent MD simulations (100 ns each). H. rPrdx6 active site’s SASA values, as a function of time, along the MD simulations at pH = 7 and at pH = 5. Data represents the average values from three independent MD simulations (100 ns each). I. Violin plots showing the distributions of Cys-47 SASA values among the MD simulations at pH = 7 and at pH = 5. Data represents the values from three independent MD simulations (100 ns each). J. Cys-47 SASA values, as a function of time, along the MD simulations at pH = 7 and at pH = 5. Data represents the average values from three independent MD simulations (100 ns each).

**Figure S5.**
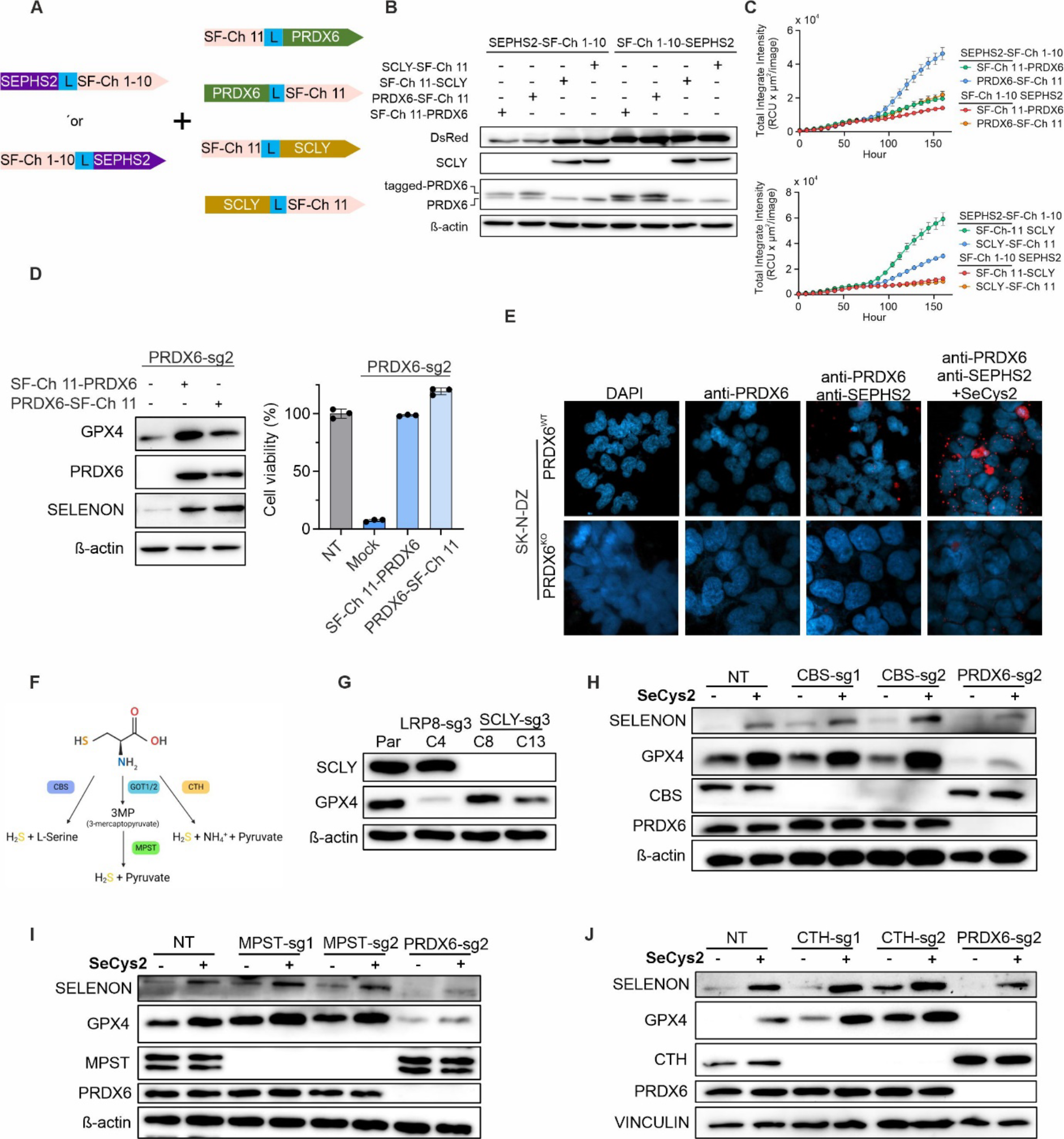
A. Schematic representation of all tested split fluorescent protein constructs used to assess the interaction of SEPHS2 with PRDX6 or SCLY. B. Immunoblot analysis of the tested split SF-Ch constructs shown in A, demonstrating the expression of the indicated proteins tagged with split SF-Ch fragments. C. Quantification of the fluorescence intensity from split SF-Ch constructs shown in A, labeling SEPHS2 together with PRDX6 (upper panel) or SCLY (lower panel) in time-lapse imaging experiments. D. Expression of GPX4 and SELENON is increased after transducing PRDX6-deficient cells with N-terminally or C-terminally tagged PRDX6 with SF-Ch 11 (left), validating the functionality of the constructs and their capacity to restore cell viability (right). E. Duolink PLA reaction in wild-type and *PRDX6* knock-out SK-N-DZ indicated interaction of PRDX6 and SEPHS2 (upper panel); Nuclei were stained with DAPI. F. Schematic representation of production of hydrogen sulfide; CBS: Cystathionine Beta- Synthase; GOT1/2: Glutamic-Oxaloacetic Transaminase 1/2; MPST: Mercaptopyruvate Sulfurtransferase; CTH: Cystathionine Gamma-Lyase. G. Immunoblot analysis of SCLY and GPX4 in wild-type SK-N-DZ, monoclonal LRP8 knock-out SK-N-DZ and monoclonal *SCLY* knock-out SK-N-DZ. H. Immunoblot analysis of SELENON, CBS, SCLY, PRDX6 and GPX4 in monoclonal SCLY knock-out SK-N-DZ transduced with sgRNA targeting *CBS* and *PRDX6* with supplement of 50 nM L-selenocystine. I. Immunoblot analysis of SELENON, MPST, SCLY, PRDX6 and GPX4 in monoclonal SCLY knock-out SK-N-DZ transduced with sgRNA targeting *MPST* and *PRDX6* with supplement of 50 nM L-selenocystine. J. Immunoblot analysis of SELENON, CTH, SCLY, PRDX6 and GPX4 in monoclonal SCLY knock-out SK-N-DZ transduced with sgRNA targeting *CTH* and *PRDX6* with supplement of 50 nM L-selenocystine.

**Figure S6.**
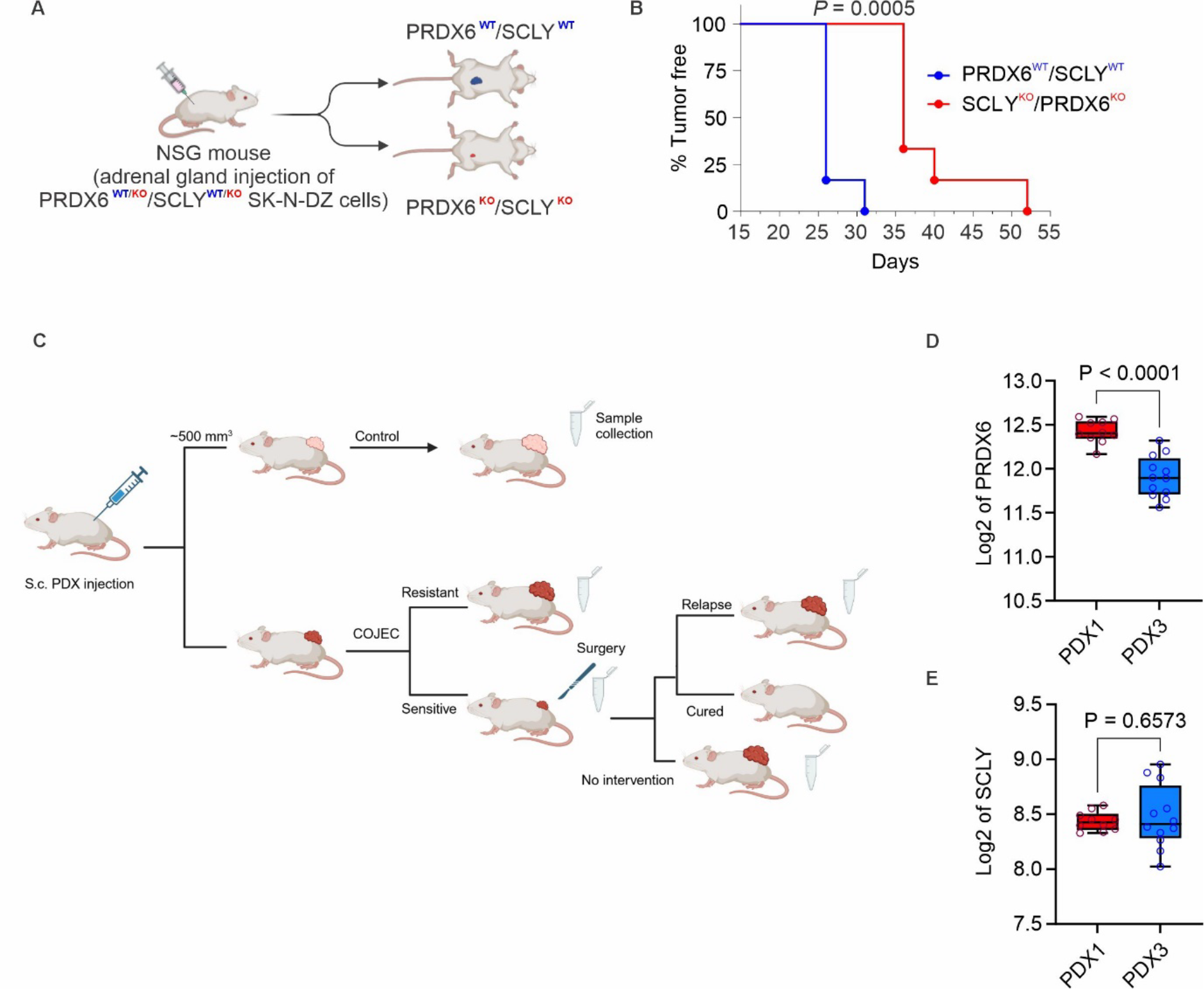
A. Schematic representation of the orthotopic implantation of control (*PRDX6^WT^/SCLY^WT^*) or PRDX6/SCLY-deficient (*PRDX6^KO^/SCLY^KO^*) SK-N-DZ cell lines. B. Kaplan–Meier plot displaying tumor-free survival (TFS) for mice injected orthotopically with *PRDX6^WT^/SCLY^WT^* (blue, n = 6) or *PRDX6^KO^/SCLY^KO^* (red, n = 6) SK-N-DZ cells. A Log-rank test was conducted for statistical analysis. C. Scheme represents the experimental design for the COJEC treatment protocol. Mice were injected subcutaneously (S.c.) with organoids from PDX1 and PDX3. After tumor achieved ∼500 mm^3^, mice were divided into groups: control and COJEC. The subgroup of PDX3, responding to COJEC treatment, was further subject to tumor resection surgery. Scalpel symbol represents surgery. Microtube symbol represents the time of collection. (The scheme is modified from Mañas et al., 2022) D. Expression of *PRDX6* in PDX1 and PDX3. E. Expression of *SCLY* in PDX1 and PDX3.

